# Intrinsic bias at non-canonical, β-arrestin-coupled seven transmembrane receptors

**DOI:** 10.1101/2021.02.02.429298

**Authors:** Shubhi Pandey, Punita Kumari, Mithu Baidya, Ryoji Kise, Yubo Cao, Hemlata Dwivedi-Agnihotri, Ramanuj Banerjee, Xaria X. Li, Cedric S. Cui, John D. Lee, Kouki Kawakami, Madhu Chaturvedi, Ashutosh Ranjan, Stéphane A. Laporte, Trent M. Woodruff, Asuka Inoue, Arun K. Shukla

**Affiliations:** Department of Biological Sciences and Bioengineering, Indian Institute of Technology, Kanpur 208016, India; Graduate School of Pharmaceutical Sciences, Tohoku University, Sendai, Japan; Department of Pharmacology and Therapeutics, McGill University, Montréal, QC, Canada; The School of Biomedical Sciences, Faculty of Medicine, The University of Queensland, Brisbane 4072, Australia; Department of Medicine, McGill University Health Center, McGill University, Montréal, QC, Canada; Queensland Brain Institute, The University of Queensland, Brisbane 4072, Australia

**Keywords:** GPCRs, biased signaling, β-arrestins, GPCR kinases, β-arrestin-coupled receptors, seven transmembrane receptors, Complement C5a

## Abstract

G protein-coupled receptors (GPCRs) are typically characterized by their seven transmembrane (7TM) architecture, and interaction with two universal signal-transducers namely, the heterotrimeric G-proteins and β-arrestins (βarrs). Synthetic ligands and receptor mutants have been designed to elicit transducer-coupling preferences and distinct downstream signaling outcomes for many GPCRs. This raises the question if some naturally-occurring 7TMRs may selectively engage one of these two signal-transducers, even in response to their endogenous agonists. Although there are scattered hints in the literature that some 7TMRs lack G-protein coupling but interact with βarrs, an in-depth understanding of their transducer-coupling preference, GRK-engagement, downstream signaling and structural mechanism remains elusive. Here, we use an array of cellular, biochemical and structural approaches to comprehensively characterize two non-canonical 7TMRs namely, the human decoy D6 receptor (D6R) and the human complement C5a receptor (C5aR2), in parallel with their canonical GPCR counterparts, CCR2 and C5aR1, respectively. We discover that D6R and C5aR2 couple exclusively to βarrs, exhibit distinct GRK-preference, and activate non-canonical downstream signaling partners. We also observe that βarrs, in complex with these receptors, adopt distinct conformations compared to their canonical GPCR counterparts despite being activated by a common natural agonist. Our study therefore establishes D6R and C5aR2 as bona-fide arrestin-coupled receptors (ACRs), and provides important insights into their regulation by GRKs and downstream signaling with direct implications for biased agonism.

## Introduction

G protein-coupled receptors (GPCRs), also referred to as seven transmembrane receptors (7TMRs), constitute a large family of cell surface proteins in the human genome with direct involvement in all major physiological processes (*1, 2*). The overall transducer-coupling framework of these receptors is highly conserved across the family where agonist-activation results in the coupling of heterotrimeric G-proteins followed by phosphorylation, primarily by GRKs, at multiple sites, and subsequent binding of β-arrestins (βarrs) (*3, 4*). While natural agonists typically induce both G-protein and βarr coupling to these receptors, it is possible to design ligands that promote preferential coupling to one of these transducers leading to biased signaling and functional outcomes (*5, 6*). This framework, referred to as biased agonism, is considered to harbor previously untapped therapeutic potential by minimizing the side-effects exerted by conventional GPCR targeting drugs (*7–9*).

A central question that still remains to be answered unequivocally is whether naturally-biased 7TMRs, able to selectively and exclusively engage one of the two well-known transducers, i.e. G-proteins and βarrs, exist? Although there are scattered examples in the literature of 7TMRs, which lack functional G-protein coupling but exhibit agonist-induced βarr recruitment (*10–12*), they are poorly characterized in terms of comprehensive G-protein coupling profile, GRK-dependence, βarr conformational signatures, and downstream signaling. These receptors include, for example, the human decoy D6 receptor (D6R) (*13–15*), the chemokine receptor CXCR7 (*16*) and the complement C5a receptor (C5L2/C5aR2) (*17–20*). Interestingly, such receptors share a natural agonist with prototypical GPCRs and therefore, constitute intriguing pairs of receptors activated by a common agonist that exhibit strikingly different transducer-coupling patterns. For example, the complement C5a peptide binds to two different 7TMRs, C5aR1 and C5aR2 but only C5aR1 exhibits functional coupling to G-protein while both of them recruit βarrs (*21, 22*).

Here, we focus on two such receptor pairs namely the CCR2-D6R activated by a common chemokine ligand CCL7, and C5aR1-C5aR2 that share complement C5a as their native agonist (Figure 1A). Using a set of complementary approaches, we discover that D6R and C5aR2 do not couple to any of the common G-proteins, robustly recruit βarrs but have differential dependence on GRKs, activate a broad spectrum of potential signaling pathways, and impart distinct conformational signatures on βarrs compared to their prototypical GPCR counterparts. This study not only establishes D6R and C5aR2 as bona fide “arrestin-coupled receptors” (ACRs) but also provides a conceptual and experimental framework that can be leveraged to discover additional receptors falling into this category, and to better understand the intricacies of biased agonism and 7TMR signaling.

**Figure 1.**
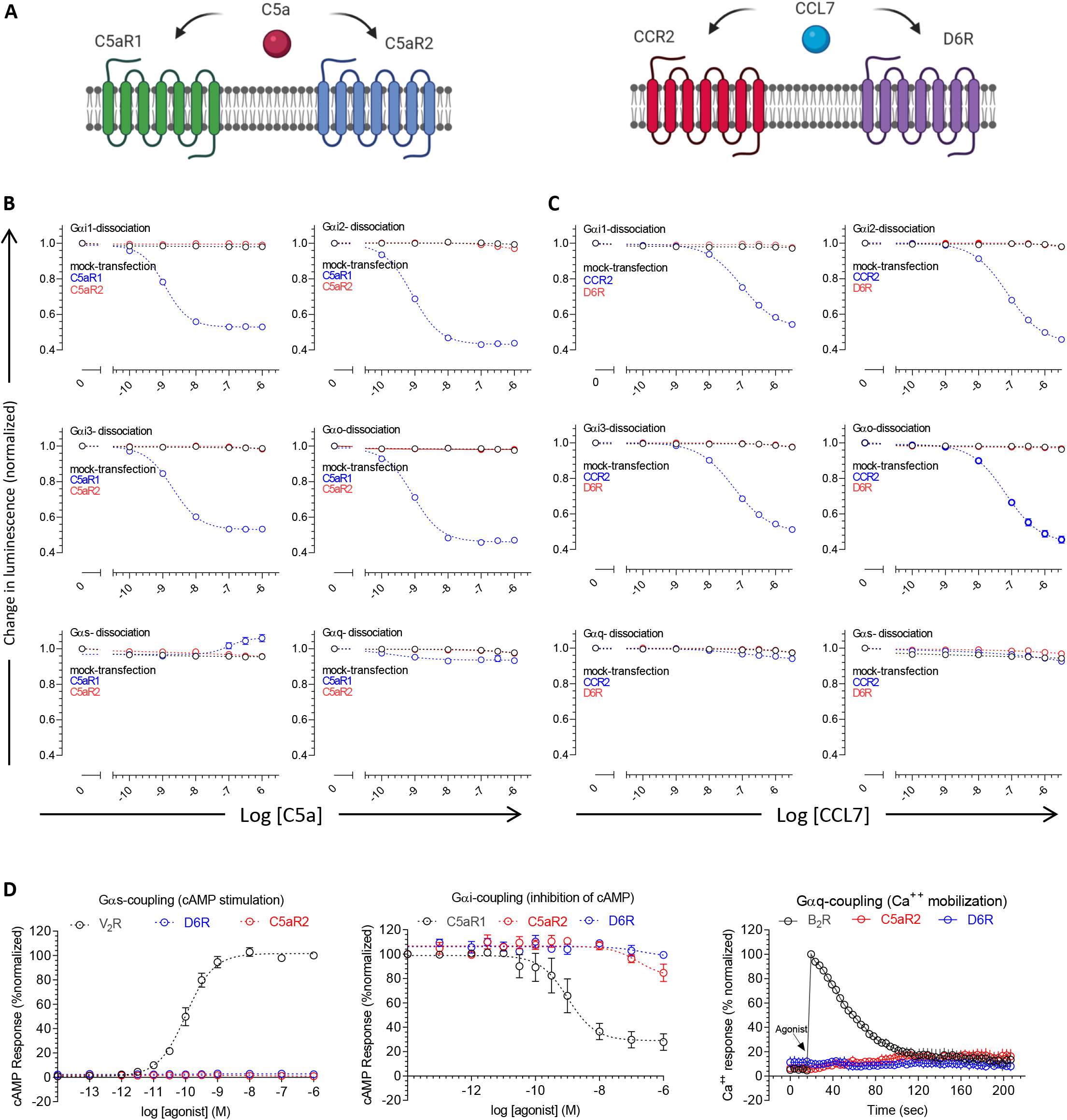
D6R and C5aR2 lack G-protein coupling and second messenger response. **A.** Schematic representation of canonical GPCR and non-canonical 7TMR pairs activated by a common agonist. C5a, complement C5a; C5aR1, C5a receptor subtype 1; C5aR2, C5a receptor subtype 2; CCL7, chemokine CCL7; CCR2, C-C chemokine receptor subtype 2; D6R, decoy D6 receptor. **B-C.** Agonist-induced dissociation of heterotrimeric G-proteins for C5aR1-C5aR2, and CCR2-D6R pairs measured using NanoBiT complementation assay. HEK-293 cells expressing the indicated receptor and Sm/Lg-BiT constructs of G-protein α,β,γ sub-units were stimulated with corresponding ligands, and the change in luminescence signal upon NanoBiT dissociation was measured as a readout of G-protein coupling. Data represent three independent experiments, normalized with respect to baseline signal (i.e. vehicle treatment). **D.** Agonist-induced second messenger response measured using GloSensor assay (cAMP stimulation, and inhibition of forskolin-induced cAMP level), and Fluo-4 NW calcium mobilization assay. HEK-293 cells expressing the indicated receptor were used for these assays using standard protocols as described in the method section. For each of the second messenger assay, a well-established prototypical GPCR was included as a reference (V_2_R, vasopressin receptor subtype 2; B_2_R, bradykinin receptor subtype 2). Data are normalized with respect to maximum signal (treated as 100%) and represent mean±sem of four independent experiments.

## Lack of G-protein coupling and second messenger response for D6R and C5aR2

Although there are scattered suggestions in the literature that D6R and C5aR2 do not couple to G-proteins, the experimental data are primarily limited to a lack of cAMP response as readout of Gαi-activation (*13, 22*). Therefore, we first set out to comprehensively measure the G-protein activation profile of these receptors using a NanoBiT-based G-protein dissociation assay (*23*). Here, a NanoBiT-G protein consisting of a large fragment (LgBiT)-containing Gα subunit and a small fragment (SmBiT)-fused Gγ2 subunit, along with the untagged Gβ1 subunit are expressed in HEK-293 cells together with the receptor construct (*23*). Subsequently, agonist-induced changes in the luminescent signal arising from the dissociation of Gα and Gβγ sub-units is measured as a readout of G-protein activation (*23*). As mentioned earlier, we used CCR2 and C5aR1 as prototypical counterparts of D6R and C5aR2 respectively, in this assay. We observed that CCR2 and C5aR1 yielded robust activation of Gαi subtype as expected, however, D6R and C5aR2 failed to generate any measurable response for any of the G-protein subtypes tested here (Figure 1B-C, Figure S1A-B). In these experiments, the receptors from each pair were expressed at comparable levels as measured by flow-cytometry-based surface expression assay (Figure S1C). Furthermore, in line with this observation, D6R and C5aR2 also failed to elicit any detectable second messenger response in cAMP and Ca^++^ mobilization assays while C5aR1 exhibited the expected profile for Gαi-coupling (Figure 1D). Taken together, these data demonstrate the lack of measurable activation of common G-proteins upon agonist-stimulation of D6R and C5aR2.

## βarr-recruitment, trafficking and GRK preference for D6R and C5aR2

In order to assess βarr-recruitment to D6R and C5aR2, we first used a co-immunoprecipitation assay by expressing these receptors in HEK-293 cells followed by agonist-stimulation, addition of purified βarrs and chemical cross-crosslinking. We observed robust interaction of βarr1 and 2 upon agonist-stimulation with each of these receptors (Figure 2A). We next monitored agonist-induced trafficking of mYFP-tagged βarr1 and 2 for D6R and C5aR2 using confocal microscopy, and we detected a typical “class B” signature of βarr trafficking as observed for other GPCRs (*24*). Upon agonist-stimulation, βarrs were first localized to the membrane followed by their trafficking to endosomal vesicles (Figure 2B-C) with an apparent preference for βarr2 (Figure 2A-C). Interestingly, we also observed some level of βarr localization to the membrane in D6R and C5aR2 expressing cells even under basal condition (i.e. before agonist-stimulation), which is more pronounced for βarr2 (Figure 2B-C). We further corroborated agonist-induced βarr1 and 2 trafficking and pre-coupling by using mCherry-tagged βarr1/2 constructs in confocal microscopy (Figure S2A-B), and scoring βarr1/2 localization patterns from a pool of cells (Figure S3).

**Figure 2.**
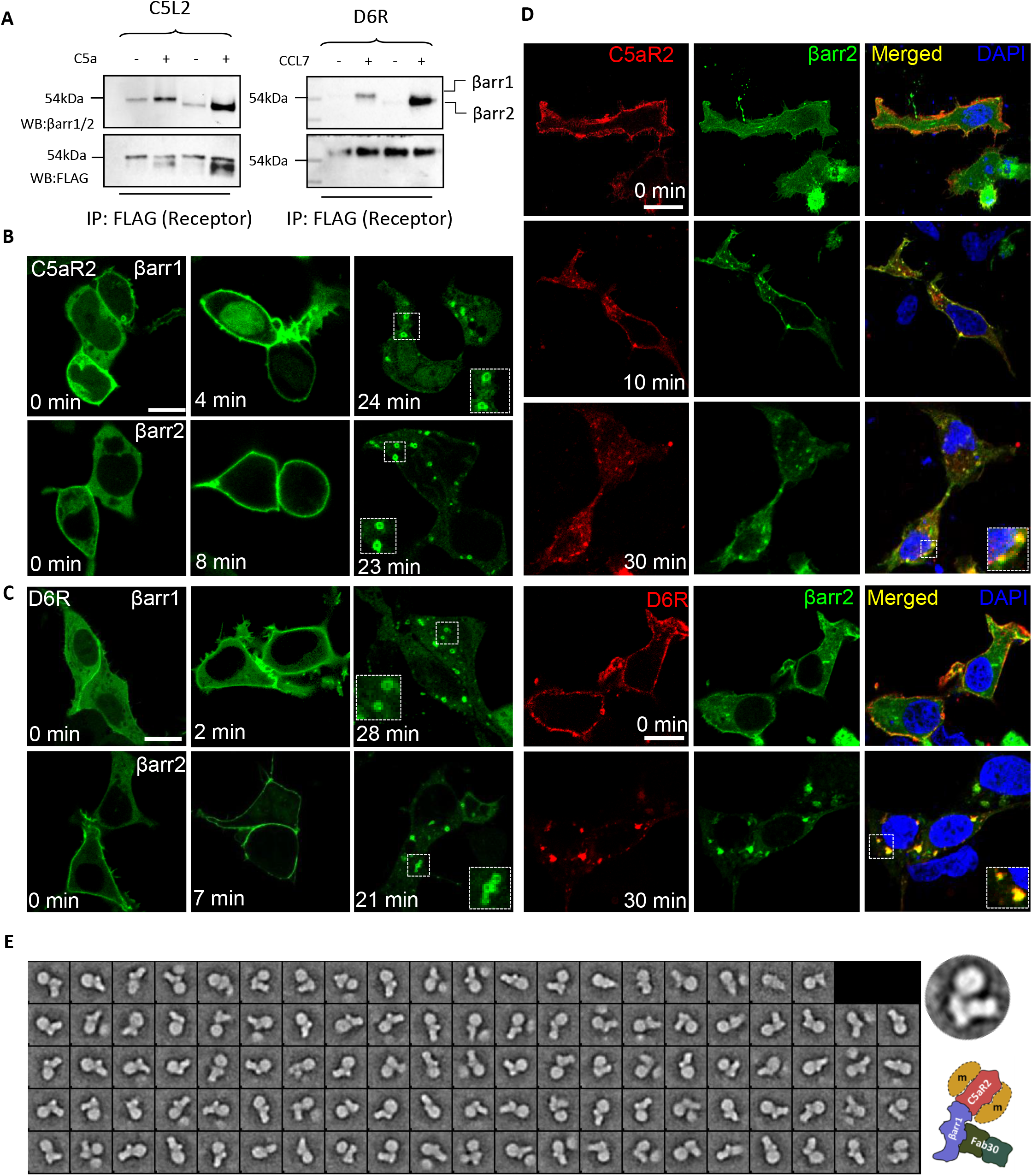
D6R and C5aR2 robustly recruit βarrs and display typical trafficking patterns. **A.** HEK-293 cells expressing the indicated receptor constructs were stimulated with agonist, lysed and incubated with purified βarr1/2. Subsequently, the receptor was immunoprecipitated using anti-FLAG M1 antibody agarose and co-purified βarrs were detected using Western blotting. A representative blot from three independent experiments is shown here. **B-C.** Agonist-induced trafficking of βarrs was measured in HEK-293 cells expressing the indicated receptor constructs and mYFP-tagged βarrs using live cell confocal microscopy. Cells were stimulated with saturating concentration of agonists (CCL7, 100 nM; C5a,100 nM) and trafficking of βarrs was monitored at indicated time-points. Representative images from three independent experiments are shown (scale bar is 10μm). **D.** Internalized D6R and C5aR2 co-localize with βarr2 as monitored by confocal microscopy. HEK-293 cells expressing the indicated receptor and βarr2-mYFP were stimulated with agonist, followed by fixation, permeabilization and staining of the receptor using DyLight-688 conjugated anti-FLAG M1 antibody. Localization of the receptor and βarr2 was visualized using confocal microscopy, and representative images from three independent experiments are shown (scale bar is 10μm). The Pearson’s Correlation Coefficient (PCC) were 0.68±0.05, 0.72±0.06, 0.94±0.01 and 0.96±0.01 for C5aR2, 0min (12 cells), C5aR2, 10min (13 cells), D6R, 0min (14 cells) and D6R, 10min (15 cells), respectively. **E.** Single particle analysis of C5aR2-V2-βarr1-Fab30 complex further corroborates the interaction of βarr1 with C5aR2. *Sf*9 cells expressing a chimeric C5aR2 construct (C5aR2-V2), GRK2^CAAX^ and βarr1 were stimulated with C5a (100 nM), stabilized using Fab30, followed by affinity purification of the complex on anti-Flag M1 column. Subsequently, the fractions containing the complex were further isolated by size-exclusion chromatography and subjected to negative staining based single particle analysis. 2D-class averages based on approximately ten thousand particles are shown here, and a typical 2D class average is indicated together with a schematic representation of the complex.

In order to probe whether internalized vesicles harbor both βarrs and receptors, we measured their colocalization by immunostaining, and observed that both D6R and C5aR2 co-localized on endosomal vesicles together with βarr2 (Figure 2D). These findings suggest that agonist-induced βarr interaction of D6R and C5aR2 has a functional consequence in terms of driving their endocytosis. In order to further establish βarr interaction, we reconstituted C5aR2-βarr1 complex stabilized by a synthetic antibody fragment (Fab30) directed against βarr1, and subjected the complex to single particle negative staining-based visualization by electron microscopy. As presented in Figure 2E, we observed several 2D class-averages reminiscent of previously described tail-engaged receptor-βarr interaction (*25*), which further confirms a direct interaction between C5aR2 and βarr1.

As receptor phosphorylation is a key determinant for βarr recruitment, and GPCR kinases (GRKs) play a central role in this process, we measured the contribution of different GRKs in agonist-induced βarr recruitment using CRISPR-CAS9 based GRK knock-out cell lines generated recently (*26*). These assays were performed under Gαi/Gαo-inhibited condition by co-expressing the catalytic subunit of pertussis toxin (PTX) in order to compare the responses for each of the receptors in the absence of G-protein signaling. We observed that C5aR2 primarily relies on GRK5/6 for βarr recruitment, a pattern that is mostly analogous to C5aR1 (Figure 3A). On the contrary, we observed that GRK knock-out has no major effect on CCL7-induced βarr recruitment for D6R, and even an increase in βarr recruitment upon GRK5/6 knock-out (Figure 3B). This is in striking contrast with CCR2, which clearly requires GRK5/6 for βarr recruitment (Figure 3B). In order to probe this interesting observation further, we assessed agonist-induced phosphorylation of D6R using the PIMAGO reagent that detects total protein phosphorylation. We observed that D6R exhibits robust constitutive phosphorylation, which does not change significantly upon CCL7-stimulation (Figure 3C). As a control, we also measured the phosphorylation of a chimeric β2-adrenergic receptor with V2R carboxyl-terminus (referred to as β2V2R), and we observed an agonist-induced increase in phosphorylation as anticipated (Figure S4A).

**Figure 3.**
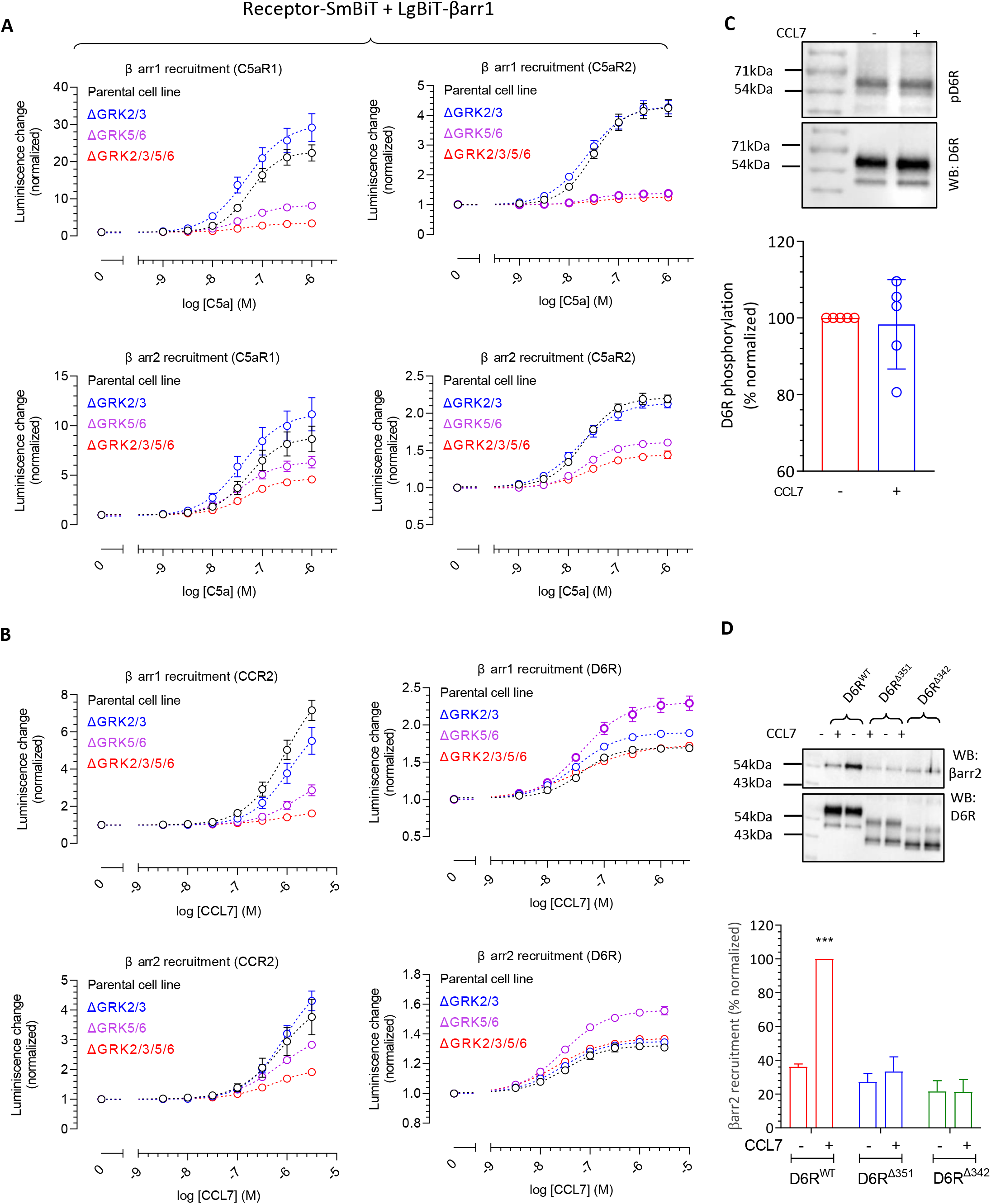
D6R and C5aR2 exhibit distinct GRK-preference for βarr recruitment. **A-B.** HEK-293 cells lacking specific GRKs were transfected with indicated receptor and βarrs followed by measurement of agonist-induced βarr recruitment using the NanoBiT assay. Data are normalized with respect to basal signal (i.e. vehicle treatment) and represent mean±sem of three independent experiments. **C.** D6R is constitutively phosphorylated as measured using PIMAGO kit, and its phosphorylation does not change upon ligand (CCL7) stimulation. HEK-293 cells expressing D6R were stimulated with CCL7 (100nM) and total receptor phosphorylation was assessed by using pIMAGO phospho-protein detection kit. Data are normalized with respect to basal signal and represent mean ± SEM of five independent experiments. **D.** HEK-293 cells expressing the indicated receptor constructs with βarr2 were stimulated with agonist followed by cross-linking and co-IP using anti-FLAG M1 antibody agarose. D6R and βarr2 were detected using Western blotting. A representative blot from three independent experiments and densitometry-based quantification is shown here. Data are normalized with D6R^WT^ stimulation condition as 100% (n=3; p<0.001).

D6R harbors a number of Ser/Thr residues in its carboxyl-terminus that represent potential phosphorylation sites (Figure S4B). In addition, it also harbors a stretch of acidic amino acids at its distal carboxyl-terminus that is suggested to play a role in its constitutive internalization (*27*). Therefore, we generated two different truncations of D6R lacking either the distal region with acidic residues (D6R^Δ351^) or the Ser/Thr cluster and the acidic residue containing stretch together (D6R^Δ342^) (Figure S4B). We observed that D6R^Δ351^ is also constitutively phosphorylated similar to D6R^WT^; however, D6R^Δ342^ did not exhibit constitutive phosphorylation (Figure S4B). These data indicate that receptor phosphorylation is localized primarily in the Ser/Thr cluster region i.e. between Ser^342^ and Ser^351^. We also measured agonist-induced βarr2 interaction and trafficking for these truncated constructs, and observed that even D6R^Δ351^ exhibits near-complete loss of βarr2 recruitment and trafficking, similar to D6R^Δ342^ (Figure 3D). These data indicate that D6R recruits βarrs primarily through the distal stretch in its carboxyl-terminus-containing acidic residues, despite having constitutive phosphorylation. In line with this observation, we also found that CCL7-stimulation fails to elicit any measurable trafficking of βarr2 for D6R^Δ351^ and D6R^Δ342^ mutants (Figure S5A-B). Taken together, these data help reconcile the intriguing observation that GRK knock-out does not influence βarr interaction for D6R.

## Distinct conformational signatures of βarrs for D6R and C5aR2

In order to probe if the distinct transducer-coupling preference of D6R and C5aR2 with respect to their prototypical GPCR counterparts may impart distinct βarr conformations, we measured the conformational signatures of βarrs upon their interaction with these receptors. First, we used a previously described intrabody30 (Ib30)-based sensor for βarr1, which selectively recognizes receptor-bound conformation of βarr1, and reports agonist-induced formation of receptor-βarr1 complex in cellular context (*28, 29*). We observed that the Ib30 sensor reacted robustly to βarr1 upon C5a-stimulation of C5aR1, but it failed to exhibit a response for C5aR2 under normalized surface expression of these receptors (Figure 4A). As C5aR2 robustly recruits βarr1, the lack of Ib30 sensor reactivity indicates a distinct conformation in C5aR2-bound βarr1 compared to C5aR1-bound βarr1. On the other hand, Ib30 recognized βarr1 for both D6R and CCR2 although the response was relatively weaker for CCR2 (Figure 4A). Considering the relatively stronger βarr1 recruitment to CCR2 thanD6R in NanoBiT assay (Figure 3B-C), it is plausible that the difference in Ib30 sensor reactivity reflects distinct conformations of βarr1 for D6R and CCR2; however, further studies are required to probe this possibility. Collectively, the Ib30 sensor data also suggests that βarr1 conformations differ between the two ACRs namely C5aR2 vs. D6R, which underscores the conformational diversity that exists in 7TMR-βarr complexes.

**Figure 4.**
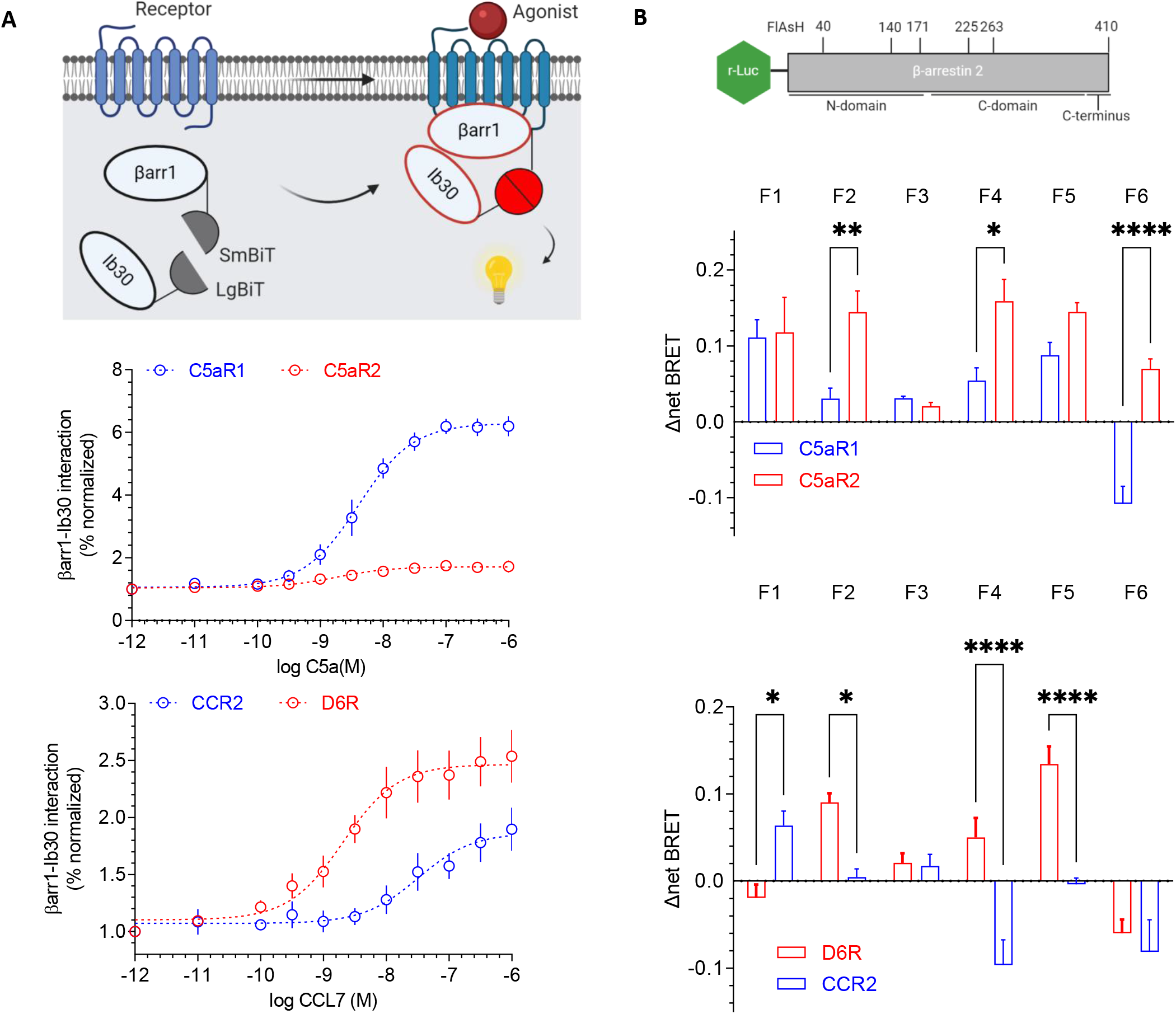
D6R and C5aR2 impart distinct conformations in βarrs compared to their GPCR counterparts. **A.** Intrabody30-based conformational sensor developed in the NanoBiT format reports the active conformation of receptor-bound βarr1 (upper panel). HEK-293 cells expressing the indicated receptor, LgBiT-Ib30 and SmBiT-βarr1 were stimulated with varying concentrations of the agonists, and luminescence signal was monitored. Data from four independent experiments (average±sem) is presented here. **B.** A set of sensors of βarr2 conformational signatures based on intramolecular BRET reveal distinct conformational signatures between C5aR1/C5aR2-βarr2 and D6R/CCR2-βarr2 complexes. The upper panel shows the schematic of BRET based conformational sensor where the N-terminus of βarr2 harbors r-Luc (Renilla luciferase) as BRET donor while FlAsH motif (as BRET acceptor) is encoded in various positions in βarr2. HEK-293 cells expressing the indicated receptor and sensor constructs were labeled with FlAsH reagent followed by agonist-stimulation and measurement of BRET signal. Data represent average±sem of four independent experiments (*p<0.05, **p<0.01, ***p<0.001).

Second, we employed FlAsH-BRET based sensors of βarr2 (*30*) to further probe the conformations of βarr2 in complex with these receptors. These intramolecular sensors harbor a BRET donor (R-luciferase) at the N-terminus of βarr2 while FlAsH labeling sequences (tetracysteine motifs) at different positions (Figure 4B). Thus, in-parallel comparison of these sensors for a given receptor can reveal conformational signatures of βarr2 with a change in BRET signal as the readout. As presented in Figure 4B, we observed striking differences not only in C5aR1-C5aR2 and D6R-CCR2 pairs but also between C5aR2 and D6R. For example, there is an opposite change in BRET signal for the F6 sensor upon activation of C5aR1 vs. C5aR2 (Figure 4B) while the F4 sensor displays directionally opposite change in BRET signal for D6R vs. CCR2 (Figure 4B). Furthermore, the comparison of BRET response for F1 and F6 sensors also reveals a distinct pattern for C5aR2 vs. D6R (Figure 4B). Taken together, these data further corroborate the conformational differences in βarr1 revealed by the Ib30 sensor, and collectively establish distinct βarr conformations induced by D6R and C5aR2 compared to their canonical GPCR counterparts. Although these assays are analytically quantitative, they are also rather qualitative in nature for assessing differences in receptor-βarr conformations, as they do not directly illuminate the precise differences unique in βarr conformations. However, they clearly reflect distinct signatures of βarrs, which can be investigated at higher resolution in the future studies using direct biophysical approaches.

## D6R and C5aR2 display distinct profile of ERK1/2 MAP kinase activation

Agonist-induced ERK1/2 MAP kinase phosphorylation has been one of the most common readouts of βarr signaling and therefore, we assessed whether D6R and C5aR2 stimulate ERK1/2 phosphorylation. While CCL7-stimulation resulted in a robust increase in ERK1/2 phosphorylation downstream of CCR2, we did not observe a detectable stimulation for D6R expressing cells (Figure 5A). We also observed a decrease in CCR2 mediated pERK1/2 at high dose of CCL7 that has been reported previously for some chemokine receptors. Interestingly, we observed a typical pattern of ERK1/2 phosphorylation upon stimulation of C5aR1; however, we noticed an elevated level of pERK1/2 in C5aR2 expressing cells, which was reduced significantly upon C5a-stimualtion in a dose dependent fashion (Figure 5B). Interestingly, the elevated level of phospho-ERK1/2 was not sensitive to pre-treatment with PTX (i.e. Gαi inhibition) (Figure 5C) but it was ablated completely with U0126 (MEK inhibitor) pre-treatment (Figure 5D). These data suggest that while a canonical pathway involving MEK is involved in the enhanced level of phospho-ERK1/2, it is not dependent on Gαi. As mentioned earlier, we observed a measurable level of constitutive βarr localization in the membrane for C5aR2 expressing cells, and therefore, we measured the effect of βarr knock-down on the basal level of ERK1/2 phosphorylation. Interestingly however, although the knock-down of βarr1 or 2 did not affect the elevated basal level of ERK1/2 phosphorylation in C5aR2 expressing cells, βarr2 depletion appears to reduce the effect of C5a on lowering ERK1/2 phosphorylation (Figure S6). Additional studies would be required to better understand the mechanistic basis of this intriguing observation in further detail.

**Figure 5.**
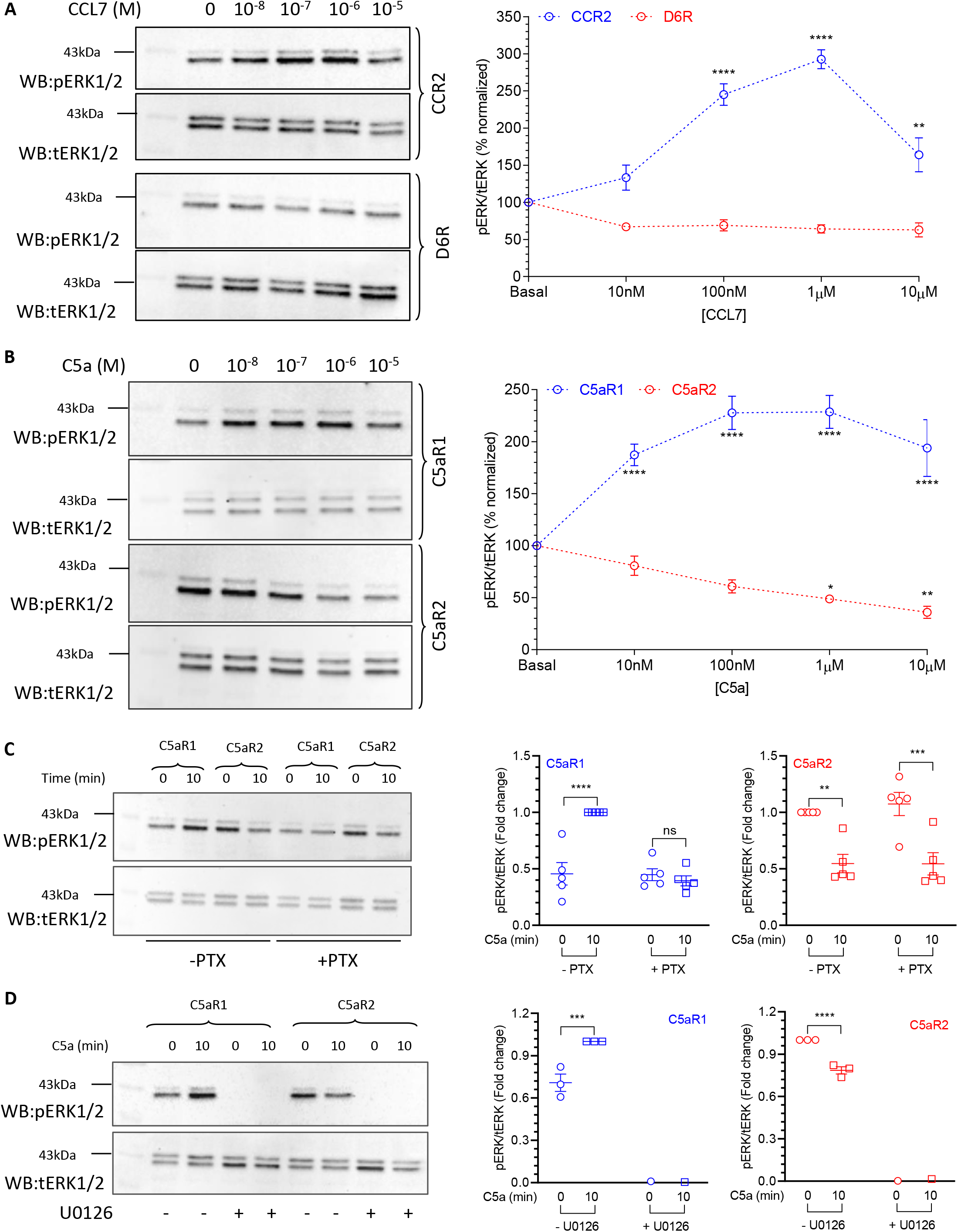
D6R and C5aR2 exhibit distinct patterns of agonist-induced ERK1/2 MAP kinase activation. **A.** CCL7 stimulation leads to robust ERK1/2 phosphorylation in HEK-293 cells expressing CCR2, however, it fails to elicit any detectable ERK1/2 phosphorylation for D6R as measured by Western blotting. **B.** C5aR1 stimulation exhibits a typical ERK1/2 phosphorylation pattern upon agonist-stimulation while C5aR2 cells display an elevated level of basal ERK1/2 phosphorylation, which decreases upon C5a-stimulation. **C.** PTX-treatment inhibits C5a-induced ERK1/2 phosphorylation downstream of C5aR1 but it fails to inhibit the elevated level of basal ERK1/2 phosphorylation for C5aR2. **D.** Treatment of cells with U0126, a MEK-inhibitor completely abolishes ERK1/2 phosphorylation for both, C5aR1 and C5aR2 suggesting the involvement of a canonical mechanism of ERK1/2 phosphorylation. The right panels show quantification based on densitometry data from 4-6 experiments analyzed using One- or Two-Way ANOVA (*p<0.05, **p<0.01, ***p<0.001, ***p<0.0001).

## Global phospho-proteomics analysis reveals potential signaling pathways downstream of D6R

In order to identify potential pathways involved in signaling downstream of D6R, we performed a phospho-antibody array based screen to identify cellular proteins that undergo a change in their phosphorylation level upon D6R stimulation. This array is designed for broad-scope protein phosphorylation profiling, and it consists of more than a thousand antibodies related to multiple signaling pathways and biological processes. In addition, we also carried out a global phospho-proteomics study on HEK-293 cells expressing D6R under basal and agonist-stimulated conditions (Figure 6A, Figure S7). These studies yielded a number of cellular proteins, which exhibit a change in their phosphorylation status upon activation of D6R (Table 1 and 2), including a sub-set that were common to both, the phospho-antibody array and phosphoproteomics (Figure S8). A classification of these proteins based on their cellular localization, molecular function and biological processes suggests that D6R activation is linked to a broad spectrum of cellular and functional outcomes (Figure 6B).

**Figure 6.**
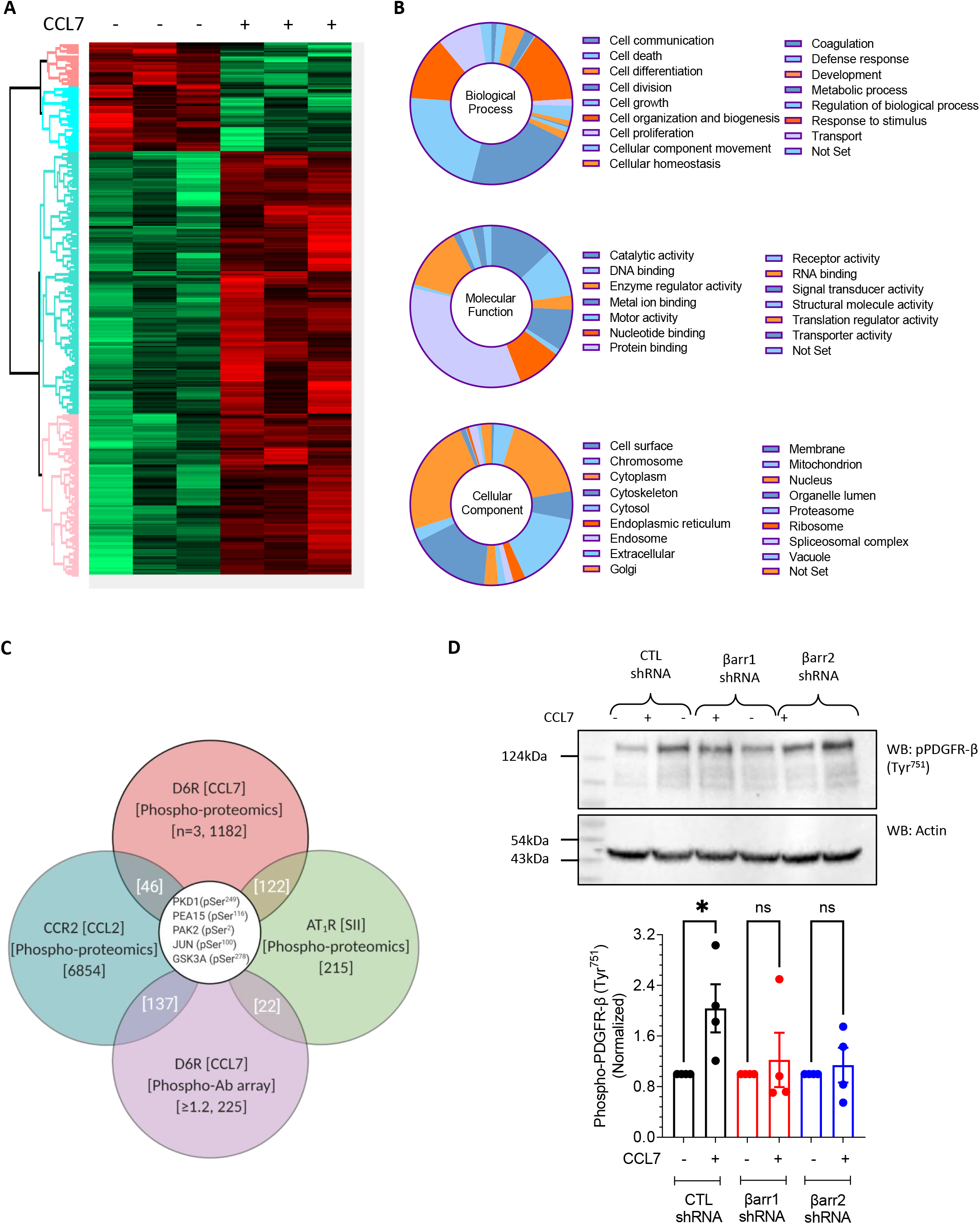
Global phosphoproteomics reveals potential signaling pathways downstream of D6R. **A.** HEK-293 cells expressing D6R were stimulated with CCL7 (100nM) followed by preparation of cellular lysate, trypsin digestion, enrichment of phospho-peptides on IMAC column and Mass-Spectrometry based identification of cellular proteins. Three independent samples prepared in parallel were analyzed and a heat map generated based on phosphoproteomics analysis is presented here. **B.** Classification of cellular proteins that undergo phosphorylation/dephosphorylation upon D6R stimulation based on biological processes, molecular functions and cellular localization reveal an extensive network of potential signaling pathways. **C.** Comparison of D6R phosphoproteomics data with phospho-antibody array and previously published βarr signaling networks reveals the activation of some common and multiple D6R-specific signaling proteins. **D.** shRNA-mediated depletion of βarr1 and βarr2 attenuate CCL7-induced (200nM) phosphorylation of PDGFR-β (Tyr^751^) in HEK-293 cells expressing D6R. A representative image from four independent experiments is shown, and the lower panel shows densitometry-based quantification, normalized with respect to the basal PDGFR-β phosphorylation (i.e. without CCL7-stimulation). Data are analyzed using One-Way ANOVA; *p<0.05).

In order to gain further insights into D6R signaling, we also compared the list of proteins identified in our study with those described previously in the context of βarr-biased agonism by stimulating the angiotensin II subtype 1 receptor (AT1R) using a βarr-biased agonist SII (*31–33*), and a recent study describing the phospho-proteome of another chemokine receptor, CCR2, in response to stimulation with CCL2 chemokine (*34*). This comparison identified not only a number of common proteins present in these studies but also several proteins that are unique to D6R activation (Figure 6C, Figure S8). These findings underline that some of the signaling downstream of D6R may be potentially similar to that identified for other GPCRs in the context of βarr-mediated pathways while there may also exist receptor-specific and previously unidentified pathways downstream of D6R. It should be noted here that SII elicits measurable Gαi and Gα12 signaling response (*35*), and therefore, a part of the hits identified earlier may not be exclusively βarr-dependent.

We also experimentally validated the phosphorylation of three different proteins namely cofilin (Ser^3^), protein kinase D (PKD) (Ser^744/748^) and the platelet-derived growth factor receptor (PDGFR-β) (Tyr^751^) upon CCL7-stimulation in D6R expressing cells, and observed agonist-induced, time-dependent response (Figure S9A-C). Furthermore, we also observed that agonist-induced phosphorylation of PDGFR-β (Tyr^751^) (Figure 6D) and cofilin (Ser^3^) (Figure S9D) is significantly attenuated upon βarr knock-down suggesting a direct involvement of βarrs. Interestingly, a previous study has reported that the phosphorylation of cofilin upon stimulation of D6R with another chemokine, CCL2, is also reduced by βarr1 depletion (*13*). Taken together these data suggest a broad signaling network downstream of D6R and set the stage for further investigation of specific signaling pathways and corresponding cellular outcomes going forward.

## C5aR2 activation leads to p90RSK phosphorylation and neutrophil mobilization

In order to probe if C5aR2 may signal through non-canonical pathways, we carried out a phospho-antibody array based screen to identify cellular proteins that undergo a change in their phosphorylation level upon C5aR1 and C5aR2 stimulation, similar to that described for D6R above. We observed that a number of proteins undergo phosphorylation/dephosphorylation upon stimulation of C5aR1 and C5aR2 expressing cells (Table1), and interestingly, several of the proteins were common to both receptors (Figure 7A) suggesting a potential involvement of βarrs. We experimentally validated agonist-induced phosphorylation of one of these proteins, p90RSK, at three different phosphorylation sites namely Thr^359^, Ser^380^ and Thr^573^. We observed agonist-induced and time-dependent phosphorylation at Ser^380^ (Figure 7B) while the other two sites did not yield consistent data. Importantly, C5a-induced phosphorylation of p90RSK at Ser^380^ is reduced upon βarr1 knock-down in HEK-293 cells suggesting an involvement of βarr1 (Figure 7C).

**Figure 7.**
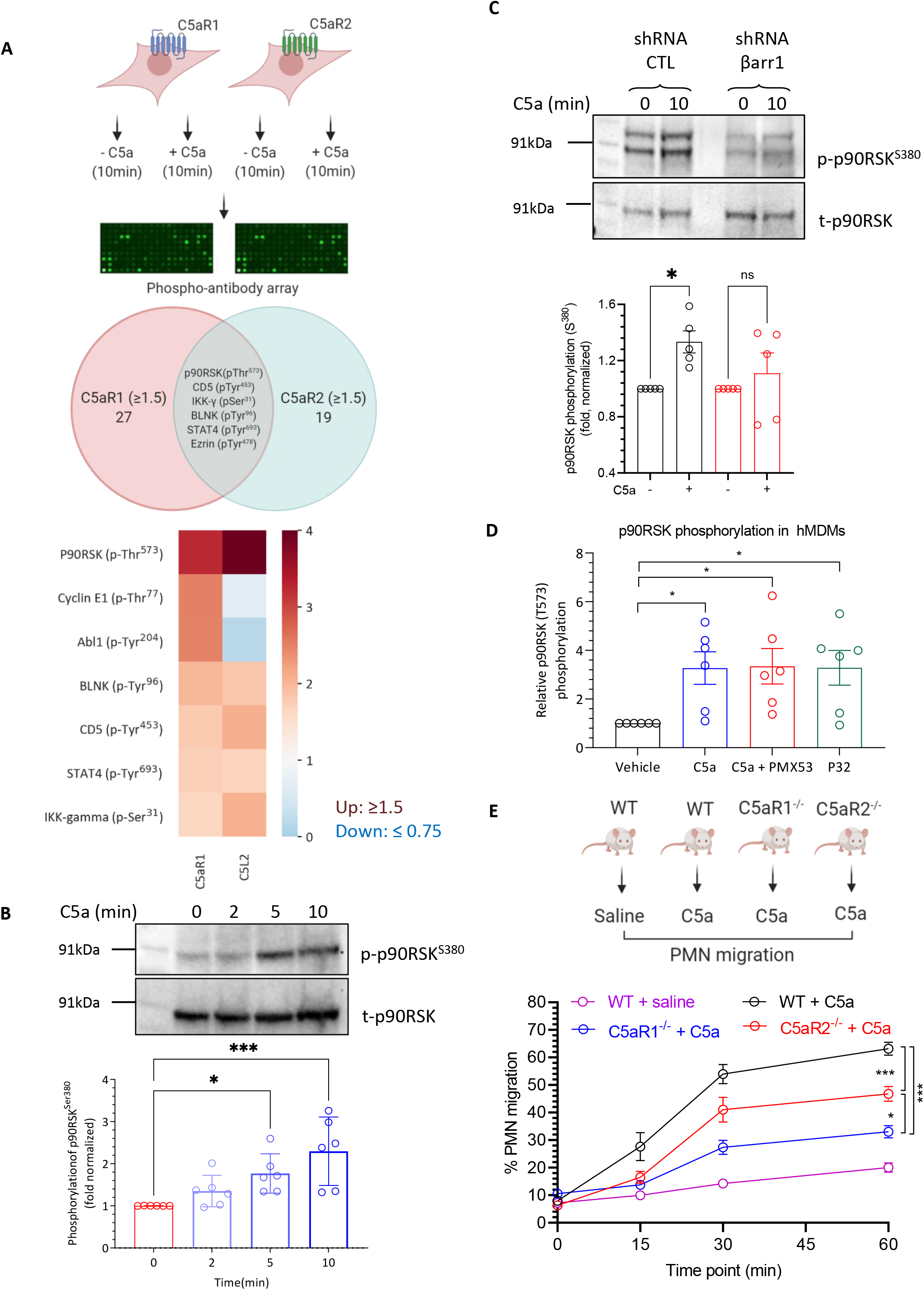
Signaling and functional outcomes of C5aR2 activation. **A.** Phospho-antibody array on HEK-293 cells expressing C5aR1 or C5aR2 reveals phosphorylation/dephosphorylation of a few common and several distinct cellular proteins. Of these, C5a-stiulation of C5aR2-expressing HEK-293 cells exhibits robust enhancement of p90RSK phosphorylation, which is also common to C5aR1. **B.** C5a-stimulation of C5aR2 expressing HEK-293 cells is validated using Western blotting, which validates the phospho-antibody array data. A representative blot from six experiments and densitometry-based quantification (average±sem) is presented (*p<0.05, ***p<0.001; One-Way ANOVA). **C.** C5a-stiulated p90RSK phosphorylation is βarr1 dependent as knock-down of βarr1 in HEK-293 cells expressing C5aR2 reduces p90RSK phosphorylation. A representative blot from five experiments and densitometry-based quantification (average±sem) is presented (**p<0.01; One-Way ANOVA). **D.** Stimulation of human macrophage derived monocytes (hMDMs) with either C5a or P32 (a C5aR2-selective agonist) results in significant p90RSK phosphorylation. Importantly, PMX53, a C5aR1-selective antagonist, does not block p90RSK phosphorylation suggesting that it is mediated by C5aR2. **E.** A significant component of C5a-induced polymorphonuclear leukocyte (PMN) mobilization depends on C5aR2. WT, C5aR1^−/−^ and C5aR2^−/−^ mice on a C57BL/6J genetic background (n = 5-15) were intravenously administered with recombinant mouse C5a (50 μg kg^−1^). Tail tip collected blood was smeared onto a slide followed by staining and counting of white blood cells, with the proportion of PMNs calculated. Data is presented as average±sem (*p<0.05, ***p<0.001; One- and Two-Way ANOVA).

In order to further corroborate this finding, we measured C5a-induced p90RSK phosphorylation in human monocyte derived macrophages (HMDMs), which constitutively express both the receptors i.e. C5aR1 and C5aR2. As efficient knock-down of βarrs in these cells has technical limitations, we used a pharmacological approach to dissect the specific contribution of C5aR2 by using a C5aR2-specific agonist (P32) (*36*). As presented in Figure 7D, we found that C5a-induced p90RSK phosphorylation at Thr^573^ in HMDMs was identical to that induced by P32. Interestingly, pre-treatment of these cells with a C5aR1 specific antagonist (PMX53) (*37*) did not block P32-induced p90RSK phosphorylation suggesting a direct involvement of C5aR2 (Figure 7D). On the other hand, C5a-induced phosphorylation of Thr^359^ and Ser^380^ in HMDMs appears to be mediated primarily by C5aR1 as pre-treatment with PMX53 blocks C5a response, and stimulation with P32 does not yield a significant response (Figure S10).

In order to identify a potential cellular and physiologically relevant effect mediated by C5aR2, we used C5aR1 and C5aR2 knock-out mice to measure C5a-induced polymorphonuclear leukocyte (PMN) mobilization. We observed that C5a induced robust PMN mobilization in wild-type mice in a time-dependent manner, which was significantly reduced but not completely abolished by C5aR1 knock-out (Figure 7E). Interestingly, PMN mobilization is also reduced in C5aR2 knock-out mice, albeit at lower levels compared to C5aR1, suggesting a distinct role of C5aR2 in PMN mobilization, in addition to the major role played by C5aR1 (Figure 7E). Taken together, these data suggest a direct contribution of C5aR2 activation in PMN mobilization and establish a distinct functional response elicited through this receptor *in vivo*.

## Discussion

The NanoBiT-based G-protein dissociation assay and second messenger assays demonstrate the lack of common G-protein coupling upon activation of D6R and C5aR2. However, an interesting question that remains to be explored is whether these receptors lack a physical interaction with G-proteins or they are incapable of activating G-proteins despite a physical interaction. In this context, it would be tantalizing to explore whether D6R and C5aR2 undergo an activation-dependent conformational change similar to that observed for prototypical GPCRs including outward movement of transmembrane (TM) helix 5 and 6 (*3*). It is plausible that activation dependent outward movement of TM5 and 6 is restricted in these receptors, which does not permit G-protein interaction; however, exploring this possibility requires additional studies. While structural elucidation of GPCR activation and signaling has seen an exponential progress, direct structural analysis of D6R and C5aR2 is still rather limited, and requires additional effort going forward.

An intriguing observation is also the constitutive phosphorylation of D6R, which does not alter significantly upon CCL7-stimulation. Although we observe some level of constitutive localization of βarrs in the membrane for D6R, it is significantly enhanced upon CCL7-stimulation followed by distribution in endosomal vesicles. Taken together with the data that βarr-recruitment is not affected upon GRK knock-out, it is possible that receptor phosphorylation has a rather minor contribution in βarr recruitment, and in fact, the truncation of distal carboxyl-terminus without disrupting the phosphorylation site cluster appears to be sufficient to ablate βarr recruitment and trafficking. These findings potentially uncover a non-canonical mode of interaction between a 7TMR and βarrs without a major role of receptor phosphorylation, which is considered as a generic paradigm in the 7TMR family.

C5aR2 has also been an enigmatic receptor since its discovery due to lack of G-protein activation, and it has been shown to be involved in broadly modulating GPCR signaling including that of C5aR1 (*21, 38*). However, the activation of a signaling pathway directly downstream of C5aR2 has not been established yet. We observe an elevated level of basal ERK1/2 phosphorylation in HEK-293 cells expressing C5aR2, which is reduced upon C5a-stimulation, but it does not appear to involve Gαi or βarrs. Importantly, we also discover that several cellular proteins undergo a change in their phosphorylation status upon C5aR2 activation, and that p90RSK phosphorylation downstream of C5aR2 is sensitive to βarr1 depletion. Therefore, our study provides a framework for exploring additional signaling pathways downstream of C5aR2, and may help uncover novel functions of C5aR2 going forward.

Finally, the distinct conformational signatures of βarrs upon their interaction with D6R and C5aR2 compared to CCR2 and C5aR1 also underscores the conformational diversity in 7TMR-βarr interaction. It is interesting to note that both D6R and C5aR2 do not exhibit the canonical readout of ERK1/2 phosphorylation despite robustly recruiting βarrs. This observation underlines the notion that recruitment of βarrs may not always translate to ERK1/2 phosphorylation, and that there may be additional cross-talk mechanisms, which fine-tune the functional responses for 7TMRs. Previous studies have linked distinct βarr conformations to different functional outcomes such as desensitization, endocytosis and signaling although a clean separation of these functional outcomes has been technically challenging (*39–43*). Our study now provides an additional handle in the form of these ACRs to decipher and link conformational signatures in βarrs to specific functional outcomes.

In summary, our study establishes D6R and C5aR2 as “arrestin-coupled receptors” with a lack of detectable G-protein coupling and potential signaling through non-canonical pathways. Moreover, we also establish that βarrs adopt distinct conformations upon interaction with these receptors compared to prototypical GPCRs, which underscores the conformational diversity of 7TMR-βarr complexes. Our findings underscore distinct functional capabilities of 7TMRs, and they have broad implications to better understand the framework of biased agonism at these receptors.

## Acknowledgement

Research in A.K.S.’s laboratory is supported by the Intermediate Fellowship of the Wellcome Trust/DBT India Alliance (IA/I/14/1/501285) awarded to A.K.S., the Swarnajayanti Fellowship of the Department of Science and Technology (DST/SJF/LSA-03/2017-18), Innovative Young Biotechnologist Award from the Department of Biotechnology (DBT) (BT/08/IYBA/2014-3), Department of Biotechnology (DBT BT/PR29041/BRB/10/1697/2018), Science and Engineering Research Board (EMR/2017/003804), Young Scientist Award from the Lady TATA Memorial Trust, and the Indian Institute of Technology, Kanpur. A.K.S. is EMBO Young Investigator, and Joy Gill Chair Professor. M.B. is supported by the National Post-Doctoral Fellowship of SERB (PDF/2016/002930) and Institute Post-Doctoral Fellowship of IIT Kanpur. H.D.-A. is supported by National Post-Doctoral Fellowship of SERB (PDF/2016/2893) and BioCare grant from DBT (BT/PR31791/BIC/101/1228/2019). M.C. is supported by a fellowship from CSIR [09/092(0976)/2017-EMR-I]. We thank Dr. Somnath Dutta for his guidance on single particle analysis using negative staining electron microscopy, Dr. Ashish Srivastava for assistance in C5aR2-βarr1 complex formation, Dr. Eshan Ghosh for helping with stable cell line generation and CCL7 purification from *Sf*9 cells, and Debarati Roy for help with CCL7 purification from *E. coli*. We thank Kayo Sato, Shigeko Nakano and Ayumi Inoue (Tohoku University) for their assistance of plasmid preparation, maintenance of cultured cells and cell-based GPCR assays. A.I. was funded by the PRIME 19gm5910013 the LEAP JP19gm0010004 from the Japan Agency for Medical Research and Development (AMED). Research performed in T.M.W’s laboratory is supported by the National Health and Medical Research Council (APP1118881). This work was also supported by a grant from the Canadian Institutes of Health Research (CIHR) (MOP‐74603) to SAL, and YC is supported by a doctoral training scholarship from the Fonds de recherche santé Québec.

## Authors’ contributions

S.P. carried out the ERK1/2 phosphorylation assays on C5aRs, p90RSK phosphorylation in HEK-293 cells, D6R phosphorylation experiments, co-IP experiments for measuring βarr interaction with D6R truncation constructs, and D6R phospho-hit validation together with H.D.-A. and M.B.; P.K. and M.B. prepared samples for phospho-antibody array experiments and phospho-proteomics, carried out data analysis and classification, βarr trafficking using confocal microscopy and ERK1/2 phosphorylation for D6R and CCR2; R.K. and K.K. carried out G-protein dissociation assays and βarr recruitment assays in GRK knock-out cells under the supervision of A.I.; Y.C. performed BRET experiments under the supervision of S.A.L.; H.D.A. performed Ib30 sensor experiments together with M.B.; R.B. carried out negative staining on C5aR2-βarr1-Fab30 complex; M.C. and A.R. assisted in validation of phospho-antibody array and phospho-proteomics hits for D6R; X.L. carried out p90RSK phosphorylation in HMDMs under the supervision of T.M.W.; S.C. and J.L. performed PMN mobilization experiment in mice under the supervision of T.M.W.; A.K.S. managed and coordinate overall project. All authors contributed to manuscript writing and editing.

## Competing interests

The authors declare that they have no competing interests.

## Data and materials availability

All data needed to evaluate the conclusions in the paper are present in the paper and the supplementary materials. Additional data related to this paper can be provided by the authors upon reasonable request.

## Materials and methods

### General reagents, plasmids, and cell culture

Most of the general chemicals and molecular biology reagents were purchased from Sigma unless mentioned otherwise. HEK-293 cells (ATCC) were maintained at 37°C under 5% CO_2_ in Dulbecco’s modified Eagle’s medium (Gibco, Cat. no. 12800-017) supplemented with 10% FBS (Gibco, Cat. no. 10270-106) and 100 U ml^−1^ penicillin and 100 μg ml^−1^ streptomycin (Gibco, Cat. no. 15140-122). Stable cell lines expressing N-terminal Flag tagged receptor constructs in pcDNA3.1 vector were generated by transfecting HEK-293 cells with 7 μg of plasmid DNA using polyethylenimine (PEI) (Polysciences, Cat. no. 19850), followed by selection using G418 (Gibco, Cat. no. 11811-031, 200-1000 μg ml^−1^). Single cell clones, which survived G418 selection, were subsequently expanded and sub-cultured. βarr1 and βarr2 shRNA expressing stable cell lines have been described earlier (*44*), and they were cultured in DMEM containing 10% (FBS), 100 U ml_-1_ penicillin and 100 μg ml^−1^ streptomycin, and 1.5μg ml-^1^ puromycin dihydrochloride (GoldBio, Cat. no. P-600). Recombinant C5a (human) and CCL7 (human) ligands were expressed and purified as described previously (*17, 45*). The plasmids encoding FLAG–C5aR1, FLAG– C5aR2, FLAG–CCR2, FLAG-D6R, βarr1–mCherry, βarr1–YFP, βarr2-YFP, Ib30-YFP have been described previously (*17, 29, 41, 44, 46*). All constructs were verified by DNA sequencing (Macrogen). The antibodies used in this study were purchased either from Sigma (HRP-coupled mouse anti-FLAG M2, HRP-coupled anti-β-actin, HRP-coupled anti-rabbit) or from Cell Signaling Technology (βarrs, ERK, PKD, PDGFRB, Cofilin, P90RSK).

### NanoBiT-G-protein dissociation assay

Agonist-induced G-protein activation was measured by a NanoBiT-G-protein dissociation assay (*23*), in which dissociation of a Gα subunit from a Gβγ subunit was monitored by a NanoBiT system (Promega). Specifically, a NanoBiT-G-protein consisting of a large fragment (LgBiT)-containing Gα subunit and a small fragment (SmBiT)-fused Gγ_2_ subunit with a C68S mutation, along with the untagged Gβ_1_ subunit, was expressed with a test GPCR construct, and the ligand-induced luminescent signal change was measured. HEK-293A cells (Thermo Fisher Scientific) were seeded in a 6-well culture plate (Greiner Bio-One) at a concentration of 2 × 10^5^ cells ml^−1^ (2 ml per dish hereafter) in DMEM (Nissui Pharmaceutical) supplemented with 10% FBS (Gibco), glutamine, penicillin, and streptomycin, one day before transfection. The transfection solution was prepared by combining 5 μl of polyethylenimine solution (1 mg ml^−1^) and a plasmid mixture consisting of 100 ng LgBiT-containing Gα, 500 ng Gβ_1_, 500 ng SmBiT-fused Gγ_2_ (C68S), and an indicated volume (below) of a test GPCR with N-terminal HA-derived signal sequence and FLAG-epitope tag followed by a flexible linker (MKTIIALSYIFCLVFADYKDDDDKGGSGGGGSGGSSSGGG; ssHA-FLAG-GPCR) in 200 μl of Opti-MEM (Thermo Fisher Scientific). To measure dissociation of the other G-protein families, LgBiT-Gα_s_ subunit (G_s_), LgBiT-Gα_q_ subunit (G_q_) and LgBiT-Gα_13_ subunit (G_13_) were used instead of the LgBiT-Gα_i1_ subunit plasmid. To enhance NanoBiT-G-protein expression for G_s_, G_q_ and G_13_, 100 ng plasmid of RIC8B (isoform 2; for G_s_) or RIC8A (isoform 2; for G_q_ and G_13_) was additionally co-transfected. To match the expression of the receptor pairs, 40 ng (C5aR1; with 160 ng of an empty vector) and 200 ng (C5aR2, CCR2 and D6R) plasmids were used. After an incubation for one day, the transfected cells were harvested with 0.5 mM EDTA-containing Dulbecco’s PBS, centrifuged, and suspended in 2 ml of Hank’s balanced saline solution (HBSS, Gibco, Cat. no. 14065-056) containing 0.01% bovine serum albumin (BSA fatty acid–free grade, SERVA) and 5 mM HEPES (pH 7.4) (assay buffer). The cell suspension was dispensed in a white 96-well plate at a volume of 80 μl per well and loaded with 20 μl of 50 μM coelenterazine (Carbosynth), diluted in the assay buffer. After 2 h incubation, the plate was measured for baseline luminescence (SpectraMax L, Molecular Devices) and 20 μl of 6X test compound (C5a or CCL7), serially diluted in the assay buffer, were manually added. The plate was immediately read for the second measurement as a kinetics mode and luminescence counts recorded from 3 min to 5 min after compound addition were averaged and normalized to the initial counts. The fold-change signals were further normalized to the vehicle-treated signal and were plotted as a G-protein dissociation response. Using the Prism 8 software (GraphPad Prism), the G-protein dissociation signals were fitted to a four-parameter sigmoidal concentration-response curve.

### Receptor surface expression assay

For measuring surface expression of the receptors, whole-cell based surface ELISA was performed as described previously (*47*). Briefly, receptor-expressing cells were seeded in 24-well plate (pre-coated with poly-D-Lysine) at a density of 0.1 million per well. After 24 h, media was removed, and cells were washed once with ice-cold 1XTBS followed by fixation with 4% PFA (w/v in 1XTBS) on ice for 20 min and subsequent extensive washing with 1XTBS. Blocking was done with 1% BSA prepared in 1XTBS for 1.5 h, which was followed by incubation of cells anti-FLAG M2-HRP antibody (Sigma, Cat no. A8592) at a dilution of 1:2000 prepared in 1% BSA+1XTBS for 1.5 h. Subsequently, cells were washed thrice with 1% BSA (in 1XTBS) to rinse off any unbound traces of antibody. Cells were incubated with 200 μL of TMB-ELISA (Thermo Scientific, Cat. no: 34028) substrate till the appearance of a light blue color and reaction was stopped by transferring 100 μL of this solution to a different 96-well plate containing 100 μL of 1 M H_2_SO_4_. Absorbance was recorded at 450 nm in a multi-mode plate reader (Victor X4, Perkin Elmer). For normalization of signal intensity, cell density was estimated using a mitochondrial stain Janus green B. Briefly, TMB was removed and cells were washed twice with 1XTBS followed by incubation with 0.2% Janus green B (Sigma, Cat. no. 201677) (w/v) for 15 min. Cells were destained by extensively washing with distilled water. The stain was eluted by adding 800 μL of 0.5 N HCl per well. 200 μL of this solution was transferred to a 96-well plate and absorbance was read at 595 nm. Data normalization was performed by calculating the ratio of A_450_ to A_595_ values.

In the NanoBiT assays, surface expression was measured using flow-cytometry based assay. Briefly, HEK-293A cells were transfected as described in the “NanoBiT-G-protein dissociation assay” section. One day after transfection, the cells were harvested with 0.53 mM EDTA-containing Dulbecco’s PBS (D-PBS). Forty percent of the cell suspension was transferred in a 96-well V-bottom plate and fluorescently labeled by using anti-FLAG epitope (DYKDDDDK) tag monoclonal antibody (Clone 1E6, FujiFilm Wako Pure Chemicals; 10 μg ml^−1^ diluted in 2% goat serum- and 2 mM EDTA-containing D-PBS (blocking buffer)) and a goat anti-mouse IgG secondary antibody conjugated with Alexa Fluor 488 (Thermo Fisher Scientific; 10 μg ml^−1^ in diluted in the blocking buffer). After washing with D-PBS, the cells were resuspended in 200 μl of 2 mM EDTA-containing D-PBS and filtered through a 40 μm filter. Fluorescent intensity of single cells was quantified by an EC800 flow cytometer equipped with a 488 nm laser (Sony). Fluorescent signal derived from Alexa Fluor 488 was recorded in a FL1 channel and flow cytometry data were analyzed by a FlowJo software (FlowJo). Live cells were gated with a forward scatter (FS-Peak-Lin) cutoff of 390 setting a gain value of 1.7. Values of mean fluorescence intensity (MFI) from approximately 20,000 cells per sample were used for analysis.

### cAMP assay

Ligand-induced Gαs- and Gαi-activation was assessed by measuring cAMP with Glosensor assay as described previously (1,4). Briefly, HEK-293 cells were transfected with FLAG-tagged receptor (3.5 μg) and luciferase-based 22F cAMP biosensor construct (3.5 μg) (Promega). 14–16 h post transfection, cells were harvested and resuspended in assay buffer containing D-luciferin (0.5 mg ml^−1^, GoldBio, Cat. no. LUCNA-1G) in 1X HBSS, pH 7.4 and 20 mM of 4-(2-hydroxyethyl)-1-piperazineethanesulfonic acid (HEPES). Cells were seeded in 96 well white plate (Corning) at a density of 125,000 cells per 100 μl and incubated at 37°C for 90 min. This was followed by an additional incubation of 30 min at room temperature. For stimulation, ligand doses (AVP for V_2_R, C5a for C5aR1 and C5aR2; CCL7 for D6R) were prepared by serial dilution ranging from 0.1 pM to 1 μM and were added to respective wells. For Gαi activation assay, prior to ligand addition, cells were treated with forskolin (5 μM). Luminescence was recorded using a microplate reader (Victor X4; Perkin Elmer). Data were normalized by treating maximal concentration of agonist as 100%. Data were plotted and analyzed using nonlinear regression in GraphPad Prism software.

### Calcium assay

In order to assess the Gαq-coupling and activation, we performed calcium assay using Fluo4-NW dye (Invitrogen, Cat. no. F36206). HEK-293 cells were transfected with FLAG-tagged receptor encoded in the pcDNA3.1 construct (3.5 μg). After 24 h of transfection, cells were seeded at a density of 125,000 cells per 50 μl in each well of a 96-well plate. Following seeding, the plate was incubated at 37°C and 5% CO_2_ for 1 h to allow the cells to settle down. After 1 h, the plate was removed from the incubator and 50 μl of freshly prepared 2X dye loading solution was subsequently added to each well. The plate was again incubated at 37°C and 5% CO_2_ for an additional 30 min followed by 30 min incubation at room temperature. The fluorescence was recorded using multimode plate reader (EnSpire; Perkin Elmer) at excitation wavelength of 494 nm and emission wavelength of 516 nm. The human bradykinin receptor (B_2_R) was used as a positive control in the experiment. Data were normalized by subtracting values of fluorescence recorded after ligand treatment with values of baseline fluorescence and time kinetics showing calcium response was plotted in the GraphPad Prism software.

### Chemical cross-linking and co-immunoprecipitation

For measuring agonist dependent βarr recruitment by respective receptor constructs, chemical crosslinking was performed following previously published protocol (5). Briefly, HEK-293 cells were transfected with FLAG-tagged receptor. 48 h post-transfection, cells were serum starved in DMEM for at least 6 h followed by stimulation with 100 nM of C5a for C5aR2 and CCL7 for D6R. Cells were lysed in a homogeniser in lysis buffer (20 mM Hepes, pH 7.4, 100 mM NaCl, Protease and Phosphatase inhibitor cocktail). Purified βarr1 or βarr2 (2.5 μg) was added to the lysate and allowed to incubate for 1 h at room temperature. Freshly prepared amine reactive crosslinker, DSP (Sigma, Cat. no. D3669) at a final concentration of 1.5 mM was added to the reaction mixture and incubated for an additional 45 min at room temperature to allow cross-linking of receptor-βarr complex. Following incubation with DSP, the reaction was quenched using 1 M Tris, pH 8.0. 1% (v/v) MNG (maltose neopentyl glycol) was added for solubilisation of receptor-βarr complex at room temperature for 1 h. In order to capture the complex, pre-equilibrated FLAG M1 antibody beads were added and incubated for additional 2 h at 4°C. Beads were thoroughly washed to remove any non-specific binding and the receptor-βarr complex were finally eluted in FLAG-EDTA solution (20 mM HEPES, 150 mM NaCl, 2 mM EDTA, 0.01% MNG, 5 μg ml^−1^ FLAG-peptide) and further incubated for 20 min at room temperature. Receptor and βarr were probed by immunoblotting by using rabbit mAb anti-βarr antibody (1:5000, CST, Cat. no. 4674). The blot was stripped and then reprobed for FLAG-tagged receptor using anti-FLAG antibody (1:2000, Sigma, Cat. no. A8592). Data were quantified using ImageLab software (Bio-Rad) and were analyzed using appropriate statistical analysis in GraphPad prism.

### Confocal microscopy

In order to visualize βarr recruitment and trafficking for D6R and C5aR2, confocal microscopy was used following the protocol described previously (*48*). Briefly, HEK-293 cells were transfected with receptor (3.5 μg), βarr1-mYFP or βarr1-mYFP (3.5 μg). For D6R mutants the constructs were normalized for surface expression and HEK-293 cells were transfected with D6R^WT^ (200 ng), D6R^1-351^ (5 μg) and D6R^1-342^ (7 μg). After 24 h, cells were seeded at 1 million density on to 0.01% poly-D-lysine (Sigma, Cat. no. P0899) pre-treated confocal dishes (SPL Lifesciences, Cat. no. 100350). Cells were allowed to attach to the plate for 24 h, prior agonist stimulation cells were serum starved for at least 6 h. For fixed cell imaging cells were fixed after starvation with 4% PFA (Sigma, Cat. no. P6148) in 1X phosphate-buffered saline (PBS, Sigma, Cat. no. D1283) for 20 min at 4°C. Fixed cells were thoroughly washed with 1XPBS and then the cells were incubated in 3% BSA with 0.1% Triton X-100 in 1XPBS (Sigma, Cat. no. 9002-93-1) for permeabilization. For staining FLAG tagged receptors we used anti-FLAG M1 antibody (1:100) labelled with DyLight594 (Thermo Scientific, Cat. no. 46412) dye in presence of 2 mM CaCl_2_. Cells were thoroughly washed with 1XPBS having 2 mM CaCl_2_. Finally, for nuclear staining, DAPI stain (5 μg ml^−1^ (Sigma, Cat. no. D9542)) was used for 10 min at room temperature. After extensive washing, cells on cover slips were mounted on slides with VectaShield HardSet mounting medium (VectaShield, Cat. no. H-1400). Confocal imaging of all samples was done using Zeiss LSM 710 NLO confocal microscope where samples were housed on a motorized XY stage with a CO_2_ enclosure and a temperature-controlled platform equipped with 32x array GaAsP descanned detector (Zeiss). A Ti: sapphire laser (Coherent) was used for exciting the DAPI channel, a Multi-Line argon laser source is used for the green channel (mYFP), and for the red channel (DyLight 594), a diode pump solid state laser source was used. All microscopic setting including laser intensity and pinhole opening were kept in the same range for a parallel set of experiments. For avoiding any spectral overlap between two channels filter excitation regions and bandwidths were adjusted accordingly. Images were scanned in line scan mode and acquired images were processed post imaging in ZEN lite (ZEN-blue/ZEN-black) software suite from ZEISS. For quantifying receptor co-localization with βarrs, the Pearson’s correlation coefficient was measured using JACoP plugin in ImageJ suite. Three regions of interest per cell were analysed for each receptor at membrane and endosomes both and the means ± S.E. of PCCs are mentioned for respective receptors in the figure legends along with the number of cells and the number of independent experiments. For quantifying βarr trafficking confocal images captured in 1 to 8 min and 9 to 30 min after agonist stimulation were categorized into early and late time points, respectively. The scoring of βarr localization was done on the basis of mYFP fluorescence either in the plasma membrane (surface localized) or in the punctate structures in the cytoplasm (internalized). In cells where βarrs were seen in both, the membrane and in punctate structures, cells having more than three punctae in the cytoplasm were scored under internalized category. All the experiments were repeated at least three times independently on different days, and the data are plotted as percentage of βarr localization from more than 500 cells for each condition. To avoid any sort of bias in manual counting, the same set of images was analyzed by three different individuals in a blinded fashion and cross-checked. All data were plotted in GraphPad Prism software.

### Isolation of C5aR-βarr1-Fab30 complex and negative staining electron microscopy

N-terminal Flag-tagged chimeric C5aR2 with V_2_R carboxyl terminus was expressed in cultured *Sf*9 cells together with GRK2^CAAX^ and βarr1 using the baculovirus expression system. 66 h post-infection, cells were stimulated with C5a followed by stabilization of the complex using Fab30. Subsequently, the complex was purified using anti-Flag M1 antibody agarose and size exclusion chromatography as described previously for the analogous β_2_AR-βarr1 complex (*25*). Prior to staining, the C5aR2-V2-βarr1-Fab30 protein complex was diluted to 0.02 mg ml^−1^ in buffer containing 20 mM HEPES, pH7.4 and 150 mM NaCl. Negative staining was performed in accordance with previously published protocols (*49*). In brief, 3.5 μl of the protein complex was applied onto glow discharged formvar/carbon coated 300 mesh copper grids (PELCO, Ted Pella) and blotted off after adsorption of the sample for 1 min using a filter paper. Staining was done with freshly prepared 0.75% (w/v) uranyl formate stain for 45 seconds. The negatively stained samples were imaged with a FEI Tecnai G2 12 Twin TEM (LaB6, 120 kV) equipped with a Gatan CCD camera (4k x 4k) at 30,000x magnifications. Approximately, 10,000 particles were picked manually from 118 micrographs using e2boxer.py within the EMAN2.31 software suite (*50*). 2D classification of the picked particles was performed with ISAC2 (*51*) within the SPHIRE suite (*52*) using the box files generated from EMAN2.31.

### NanoBiT-β-arrestin recruitment assay

The effect of GRKs on agonist-induced βarr activation was measured by a NanoBiT-βarr recruitment assay. Specifically, the parental HEK-293A, GRK2/3-KO, GRK5/6 and GRK2/3/5/6-KO cells (*26*) were seeded and transfection was performed by following the same procedures as described in “NanoBiT-G-protein dissociation assay” section. For the βarr recruitment assay, 100 ng (hereafter per well in a 6-well plate) of an N-terminally LgBiT-fused βarr1/2 plasmid and 80 ng (C5aR1; with 420 ng of an empty vector) or 500 ng (C5aR2, CCR2 and D6R) of a test GPCR plasmid with the N-terminal HA-derived signal sequence and FLAG-epitope tag and a C-terminal SmBiT (ssHA-FLAG-GPCR-SmBiT). The transfected cells were subjected to the NanoBiT luminescent measurement as described above. Luminescence counts from 10 min to 15 min after compound addition were used for the calculation.

### Detection of D6R basal phosphorylation by pIMAGO assay

To detect agonist independent basal phosphorylation in D6R, pIMAGO phosphoprotein detection kit from Sigma (Cat. no. 18419) was used and receptor phosphorylation was detected as per the manufacturer’s protocol. Briefly, HEK-293 cells were transfected with 7 μg D6 receptor DNA complexed with 21 μg PEI. 48 h after transfection, cells were serum-starved for 6 h followed by stimulation with 200 nM CCL7 for 30 min and harvested in 1XPBS. Post stimulation, cells were lysed in buffer containing 50 mM HEPES (pH 7.4), 150 mM NaCl, 10% glycerol (v/v), 1% NP40, 2 mM EDTA, 1X phosSTOP and 1X protease inhibitor cocktail (Roche, Cat. no. 04693116001) for 2h at room temperature. The lysate was cleared by centrifugation and transferred to a separate tube already containing pre-equilibrated M1-FLAG beads supplemented with 2 mM CaCl_2_. The receptor was enriched by performing FLAG-immunoprecipitation as described previously. Afterwards, the protein was eluted in FLAG elution buffer containing 20 mM HEPES pH 7.5, 150 mM NaCl, 2 mM EDTA, 0.06% NP40 and 250 μg ml^−1^ FLAG peptide. Subsequently, protein loading dye was added to each sample, followed by the addition of 5X IAA solution to a 1X final concentration from the pIMAGO kit. The samples were incubated at room temperature for 15 min in dark. Eluted samples were subjected to SDS-PAGE followed by western blotting. The membrane was blocked in 1X blocking buffer for 1 hfollowed by incubation with pIMAGO reagent (1:1000, prepared in 1X pIMAGO buffer) for 1 h. The membrane was washed thrice with 1X wash buffer and once with 1XTBST (5 min each wash). The PVDF membrane was incubated with avidin-HRP (1:1000, prepared in 1X blocking buffer) for 1h at room temperature and washed thrice with 1XTBST (5 min each wash). The signal was detected using Promega ECL solution on chemidoc (BioRad). Blot was stripped and re-probed for total receptor using HRP conjugated anti-FLAG M2-antibody (Sigma, 1: 5000). The signal was normalized with respect to total receptor and quantified using ImageLab software (BioRad).

To assess the role of D6R C-terminus in basal receptor phosphorylation, the receptor was truncated at C-terminus at two positions (i.e. 1-342 and 1-351) by inserting a STOP codon by site-directed mutagenesis (NEB, Cat. no. E0554). The surface expression of WT and truncated receptor constructs were normalized to similar levels by DNA titration in HEK-293 cells. Relative surface expression of all the constructs was measured by whole cell-based surface ELISA as described previously. For the detection of basal phosphorylation in D6R-WT and mutants, 50-60% confluent HEK-293 cells were transfected with D6R-WT (200 ng) and mutant receptor DNA complexed with 21 μg PEI (1-342: 7 μg, 1-351: 5 μg). For each construct, 5×10 cm HEK-293 plates were transfected. 48 h post-transfection, cells were harvested in 1XPBS and lysed in NP40-lysis buffer. Receptor phosphorylation was detected using a western blot based pIMAGO-phosphoprotein detection kit as mentioned in the previous section.

To identify the specific determinants of βarr interaction in D6R C-terminus, 50-60% confluent HEK-293 cells were transfected with either D6R-WT (200 ng) or mutants (1-342: 5 μg and 1-351: 5 μg) and βarr2 (2 μg). The surface expression of WT and mutant receptor constructs was normalized to similar levels as mentioned in the previous section. 48h post-transfection, cells were serum-starved for 6 h followed by stimulation with 100 nM CCL7 for 30 min. Post-stimulation, cells were harvested in 1XPBS and proceeded for chemical crosslinking. Cells were lysed by Dounce homogenization in 20 mM HEPES pH 7.5, 350 mM NaCl, 1XPhosSTOP, and 1X complete protease inhibitor cocktail). This was followed by the addition of freshly prepared dithiobis(succinimidyl-propionate) to a final concentration of 1 mM. Cell lysates were tumbled at room temperature for 40 min and the reaction was quenched by 1 M Tris pH 8.5. Afterward, lysates were solubilized in 1% MNG (w/v) at room temperature for 1.5 h and centrifuged at 15000 rpm for 10 min. Cleared lysates were supplemented with CaCl_2_ to a final concentration of 2 mM followed by the addition of pre-equilibrated M1-FLAG beads to the lysate. The samples were tumbled at room temperature for 1.5 h to allow bead binding and beads were washed 3 times each with low salt buffer (20 mM HEPES pH7.5, 150 mM NaCl, 2 mM CaCl_2_, and 0.01% MNG) and high salt buffer (20 mM HEPES pH7.5, 350 mM NaCl, 2 mM CaCl_2_ and 0.01% MNG) alternately. The bound proteins were eluted in FLAG-elution buffer containing 20mM HEPES pH 7.5, 150mM NaCl, 2mM EDTA, 0.01% MNG and 250 μg ml^−1^ FLAG peptide. Eluted βarr2 was detected by Western blotting using rabbit anti-βarr mAb (1:5000, CST, Cat. no. 4674). The blots were stripped and re-probed for receptor with HRP-coupled anti-FLAG M2 antibody (1:5000). The blots were developed on Chemidoc (Bio-Rad) and quantified using ImageLab software (Bio-Rad).

### Ib30 NanoBiT Assay

We measured ligand-induced βarr conformational change recognized by Intrabody 30 (Ib30) using NanoBiT assay (*28*). Ib30 and βarr1 were N-terminally fused to LgBiT and SmBiT respectively with the 15-amino acid flexible linker and inserted into the pCAGGS plasmid. The receptor pair C5aR1 and C5aR2 exhibited matched cell surface expression at DNA concentration of 0.25 μg and 3 μg respectively. Similarly, cells transfected with 0.5 μg DNA of D6R and 3 μg CCR2 showed comparable surface expression. For NanoBiT assay, HEK-293 cells at a density of 3 million were transfected with receptor (DNA concentration as mentioned above), LgBiT-Ib30 (5 μg) and SmBiT βarr1 (2 μg) using PEI (Polyethylenimine; 1 mg ml^−1^) as transfection agent at DNA:PEI ratio of 1:3. After 16-18 h of transfection, cells were harvested in PBS solution containing 0.5 mM EDTA and centrifuged. Cells were resuspended in 3 ml assay buffer (HBSS buffer with 0.01% BSA and 5 mM HEPES, pH 7.4) containing 10 μM coelenterazine (Goldbio, Cat. no: CZ05) at final concentration. The cells were then seeded in a white, clear-bottom, 96 well plate at a density of 0.7 × 10^5^ cells per100 μl per well. The plate was kept at 37:C for 90 min in the CO_2_ incubator followed by incubation at room temperature for 30min. Basal reading was read on luminescence mode of multi-plate reader (Victor X4). The cells were then stimulated with varying doses of each ligand (C5a and CCL7) ranging from 0.1 pM to 1 μM (6x stock, 20 μl per well) prepared in drug buffer (HBSS buffer with 5 mM HEPES, pH 7.4). Luminescence was recorded for 60 min immediately after addition of ligand. The initial counts of 4-10 cycles were averaged and basal corrected. Fold increase was calculated with respect to vehicle control (unstimulated values) and analyzed using nonlinear regression four-parameter sigmoidal concentration–response curve in GraphPad Prism software.

### FlAsH BRET experiments

HEK-293SL cells were cultured in DMEM supplemented with 10% FBS and 20 μg ml^−1^ gentamicin, and grown at 37°C in 5% CO_2_ and 90% humidity. Cells were seeded at a density of 1.5×10^5^ cells per well in 6-well plate and were transiently transfected the next day with C5aR1, C5aR2, D6R, or CCR2 and βarr2-FlAsH constructs using conventional calcium phosphate co-precipitation method. One day post-transfection, cells were detached and seeded in poly-ornithine-coated white 96-well plates at a density of 2.5×10^4^ cells per well in media. The next day, cells were washed and incubated for 1 h with Tyrode’s buffer (140 mM NaCl, 2.7 mM KCl, 1 mM CaCl_2_, 12 mM NaHCO_3_, 5.6 mM D-glucose, 0.5 mM MgCl_2_, 0.37 mM NaH_2_PO_4_, 25 mM HEPES, pH 7.4) at room temperature. FlAsH labeling was performed as previously described (*30*). Briefly, 1.75 μl of FlAsH-EDT_2_ stock reagent was mixed with 3.5 μl of 25 mM EDT solution in DMSO and left for 10 min at room temperature. 100 μl of Tyrode’s buffer was then added and left for 5 min at room temperature. The volume was then adjusted to 5 ml with Tyrode’s buffer to complete the labeling solution. Cells were washed with Tyrode’s buffer and incubated with 60 μl of labeling solution per well for 1 h at 37°C. Cells were then washed twice with BAL wash buffer followed by another wash with Tyrode’s buffer. Next, 90 μl of Tyrode’s buffer was added per well and incubated for 1 h at 37°C. Cells were stimulated with 1 μM C5a or CCL7 ligand for 10 min, with six consecutive BRET measurements taken every minute after 5 min stimulation. Cell-permeable substrate coelenterazine H (final concentration of 2 μM) was added 3 min prior to BRET measurements, with triplicates for each condition. BRET measurements were performed using a Victor X (PerkinElmer) plate reader with a filter set (center wavelength/band width) of 460/25 nm (donor) and 535/25 nm (acceptor). BRET ratios were determined by dividing the intensity of light emitted by the acceptor over the intensity of light emitted by the donor. The net BRET ratio is calculated by subtracting the background BRET ratio (unlabeled) from the FlAsH-EDT_2_-labeled BRET ratio. The Δnet BRET is then obtained by dividing the stimulated net BRET ratio by the vehicle net BRET ratio.

### ERK1/2 phosphorylation assay in HEK-293 cells

Agonist-induced ERK1/2 MAP kinase phosphorylation was carried out following the protocol described previously (*53*). Briefly, HEK-293 cells expressing the indicated receptors were seeded into a 6-well plate at a density of 1 million cells per well. Cells were serum-starved for 12 h followed by stimulation with the indicated concentration of corresponding ligands at specific time points. To study the effect of pertussis toxin (PTX) on basal ERK phosphorylation of C5aR2, cells were treated with 100 ng ml^−1^ PTX (in starvation media) for 12 h prior to ligand stimulation. Similarly, to study the effect of MEK-inhibitor (U0126) on basal ERK phosphorylation of C5aR2, cells were pretreated with 10 μM U0126 for 30 min before ligand stimulation. After the completion of the time course, the media was aspirated, and cells were lysed in 100 μl 2x SDS dye per well. Cell lysates were heated at 95:C for 15 min followed by centrifugation at 15000 rpm for 10 min. 10 μL of lysate was loaded per well and separated in SDS-PAGE followed by western blotting. Blots were blocked in 5% BSA (in TBST) for 1 h and incubated overnight with rabbit phospho-ERK (CST, Cat. no. 9101) primary antibody at 1:5000 dilution. Blots were washed thrice with TBST for 10 min each and incubated with anti-rabbit HRP-coupled secondary antibody (1:10000, Genscript), Cat. No. A00098 for 1 h. Blots were washed again with TBST for three times and developed with Promega ECL solution (Cat. no. W1015) on chemidoc (BioRad). Blots were stripped with low pH stripping buffer and then re-probed for total ERK using rabbit total ERK (CST, Cat. no. 9102/) primary antibody at 1:5000 dilution.

### Phospho-antibody array

A phospho-antibody array (Full Moon Biosystems) consisting of 1318 antibodies against proteins from multiple signaling pathways were used to discover potential signaling pathways downstream of receptors investigated here. The samples were prepared as per the manufacturer’s instruction and sent to Full Moon Biosystems for further analysis. Briefly, HEK-293 cells stably expressing the receptor was stimulated with saturating concentration of ligands (C5a, 100 nM for C5aR1 and C5aR2; CCL7, 100 nM for D6R) for 10 min and then harvested using 1 ml of ice-cold 1X PBS supplemented with 0.01% Phosphatase inhibitor (PhosSTOP, Roche, Cat. no. 04906845001). Pellets corresponding to 10 plates of a 10 cm plate were pooled together and centrifuged at 5,000 rpm for 5 min at 4°C. The supernatant was discarded and the pellets were washed again with 1 ml of cold 1X PBS to remove any traces of media. HEK-293 cells stably expressing the receptor under non-stimulation conditions were used as a control. Three independent set of pellets comprising of cells pooled from the unstimulated conditions and stimulated conditions were prepared following similar conditions and sent to Full Moon Biosystems for phosphoarray and analysis. The antibody array was done using a kit (Cat. no. KAS02). Briefly cells was lysed and centrifuged to obtain a clear lysate. Prior labelling of the proteins in the lysate with biotin, buffer was exchanged for ensuring proper biotinylation. The amount of total protein was analyzed using BCA estimation for both unstimulated and stimulated conditions. Subsequently, equal amount of biotinylated proteins were allowed to bind with the immobilized antibodies coated on a glass slide. After rigorous washing, Cy3-streptavidin was used to detect the bound proteins to respective antibodies. The antibody array slide was finally detected using a microarray scanner. Fold increase in signal was obtained after dividing the fluorescence signal emitted from respective antibody spots for stimulated sample by corresponding signal from unstimulated sample.

### MS-based Phosphoproteomics

For MS-based phosphoproteomics of D6R, HEK-293 cells stably expressing the D6R were grown at a confluency of ~70%. Cells were serum starved for at least 6 h prior to stimulation. Cells were then stimulated for 10 min with 100 nM of CCL7. Media was then aspirated and cells were washed with 1XTBS containing 0.01% of phosphatase inhibitors. Cells corresponding to 10 plates each were scraped and collected and pelleted in a 15 ml falcon. Three independent sets of unstimulated and stimulated cell pellets each were prepared. The pellets were then treated with 6 M Gn-HCL/0.1 M Tris (pH 8.5) plus phosphatase inhibitors and resuspended well. The lysate was then boiled at 90°C for 10 min, followed by sonication for breaking the nucleic acids and reducing the viscosity of slimy material. After sonication, lysate was again boiled at 90°C for 5 min and then spun at 15,000 rpm for 20 min at room temperature. The supernatant was collected carefully leaving behind the cell pellet. Protein concentration was estimated by BCA using the same lysis buffer as a blank solution. Lysates corresponding to 5 mg or more was transferred into 1.5 ml Eppendorf tubes, double parafilmed and sent to V-Proteomics for analysis.

For sample preparation, 25 μg of the sample from each condition were first reduced with 5 mM TCEP and further alkylated with 50 mM iodoacetamide. Alkylated proteins were then digested with trypsin (1:50, trypsin: lysate ratio) for 16 h at 37°C. Prior to phospho-enrichment of the samples with TiO_2,_ digested samples were cleaned up using Sep-Pak columns. These digests and enriched samples were further cleaned up using C18 silica cartridge and dried using speed vac. The dried pellet was resuspended in Buffer-A (5 % acetonitrile / 0.1% formic acid). For the mass spectrometric analysis of peptide mixtures, all the experiments were performed using EASY-nLC 1000 system (Thermo Fisher Scientific) coupled to QExactive mass spectrometer (Thermo Fisher Scientific) equipped with nanoelectrospray ion source. 1 μg of the peptide mixture was loaded on pre-column and resolved using 15 cm Pico-Frit filled with 1.8 μm C18-resin (Dr. Maeisch). The peptides were loaded with Buffer A and eluted with a 0-40% gradient of Buffer-B (95% acetonitrile/0.1% Formic acid) at a flow rate of 300 nl min^− 1^ for 105 min. The QExactive was operated using the Top10 HCD data-dependent acquisition mode with a full scan resolution of 70,000 at m/z 400. MS/MS scans were acquired at a resolution of 17500 at m/z 400. Lock mass option was enabled for polydimethylcyclosiloxane (PCM) ions (m/z = 445.120025) for internal recalibration during the run. MS data was acquired using a data-dependent top10 method dynamically choosing the most abundant precursor ions from the survey scan.

For data analysis, all six raw files (3 sets of stimulated and 3 sets of unstimulated samples) were analyzed with Proteome Discoverer 2.2 against the Uniprot Human reference proteome database (containing 20162 entries). For Sequest HT and MS Amanda 2.0 search, the precursor and fragment mass tolerances were set at 10 ppm and 0.5 Da, respectively. The protease used to generate peptides, i.e. enzyme specificity was set for trypsin/P (cleavage at the C terminus of “K/R: unless followed by “P”) along with maximum missed cleavages value of two. Carbamidomethyl on cysteine as fixed modification and oxidation of methionine and N-terminal acetylation were considered as variable modifications for database search. Both peptide spectrum match and protein false discovery rate were set to 0.01 FDR and determined using percolator node. Relative protein quantification of the proteins were performed using Minora feature detector node of Proteome Discoverer 2.2 with default settings and considering only high PSM (peptide spectrum matches) confidence. Based on uniprot accession number Pfam, KEGG pathways and GO annotations were assigned for the list of identified proteins. Also, for high sensitivity-phospho site localization to be detected for individual site, ptmRS node was considered. Based on relative abundance of phospho-peptides across three independent sets a one-way ANOVA test was performed to screen out significant and high confidence phospho-peptides by using Perseus (MaxQuant).

In order to get a functional insight from the robust data generated from phospho-proteomics and phospho-array, we also analyzed additional data sets from the literature namely the phospho-proteomics data set for CCR2, a chemokine receptor (*34*), and a βarr biased phospho-proteomics data set carried out on the AT1aR stimulated with a biased agonist SII (*31, 33*). The rationale behind including the above datasets is because D6R is a chemokine receptor with exclusive bias towards βarr. Therefore, comparing the above-mentioned dataset can bring out some common features of βarr signaling involving chemokine receptors. Moreover, the unique hits from D6R not matching with these datasets will allow us to understand novel βarr mediated signaling outcome in atypical chemokine receptors. After analyzing all data sets overlapping hits were identified. The phosphor-sites mentioned for the common identified proteins from all the data sets in Figure 6C are retrieved from D6R phosphor-proteomics data set. Furthermore, a list of unique proteins was generated from both phosphor-array and phosphor-proteomics data of D6R and the unique proteins with their phospho-sites labeled were submitted to kinase enrichment analysis tool KEA 2.0 (https://www.maayanlab.net/KEA2/). The significantly enriched hits were listed based on their P-values. The listed proteins were further analyzed in STRING database to identify protein-protein interaction to understand the pathways involved. Using this criterion, we selected three proteins i.e. protein kinase D1 (PKD1), cofilin, and the platelet derived growth factor receptor β (PGGFR-β) for immunoblotting based validation from cell lysate.

### Validation of D6R phospho-protein hits

For validation of phospho-array and phospho-proteomics hits, stable cell lines expressing D6R were seeded at 3 million per 10 cm plate. Cells were serum-starved with 20 mM HEPES (pH 7.4) and 1% BSA in serum-free DMEM media for 16-18h. Cells were then stimulated with 200 nM CCL7/for indicated time points. Cells were lysed in buffer containing 50 mM HEPES (pH 7.4), 150 mM NaCl, 10% glycerol (v/v), 1% NP40, 2 mM EDTA, 1X phosSTOP and 1X protease inhibitor cocktail for 1h at room temperature. The lysate was cleared by centrifugation and solubilized proteins were estimated with BCA method (G Bioscience). Approximately 90-100 μg of each sample was loaded on 4-20% precast gradient gel (Bio-Rad) and resolved proteins were transferred to a PVDF membrane. After blocking with 3% BSA in 1X TBST blots were incubated with primary antibody (phospho-Cofilin^S3^, 1:1000; phospho-PKD^S744/748^1:1000, phospho-PDGFR^Y751^, 1:1000). Blots were developed in ChemiDoc™MP Gel Imaging System (Bio-Rad) after incubating the blots in anti-rabbit HRP antibody (1:5000) for 1h at room temperature.

To evaluate the role of βarr isoforms in phosphorylation of these hits, D6R or C5aR2 plasmids were transfected in control-, βarr1- and βarr2-shRNA expressing cell lines at 7 μg. Serum-starvation, stimulation, and sample preparation were performed as mentioned previously. About 90-100 μg of cell lysates were run on 4-20% precast gradient gel and western blotting was performed as per the previous protocol. PVDF membranes were probed for phospho-Cofilin^S3^ (CST, Cat. no. 3313, 1:1000), phospho-PKD^S744/748^ (CST, Cat. no. 2054, 1:1000), phospho-PDGFR^Y751^(CST, Cat. no. 4549, 1: 1000), phospho-P90RSK^S380^ (CST, Cat. no. 11989, 1:500). Blots were stripped with low pH stripping buffer and then re-probed for total RSK using RSK1/RSK2/RSK3 rabbit monoclonal primary antibody at 1:2500 dilution (CST, Cat. no. 9355S,) or for β-actin (Sigma, Cat. no. A3854, 1:50000). Phospho-site specific signal was normalized with respect to the total RSK or β-actin signal.

### Ligand-induced p90RSK phosphorylation in HEK-293 cells

In order to measure C5a-induced p90RSK phosphorylation, HEK-293 cells stably expressing C5aR2 were seeded in 10 cm culture dishes at a density of 5 million. After 24 h, cells were subjected to serum starvation for 16 h followed by stimulation with 100 nM C5a for indicated time points and harvested in PBS. Subsequently, cells were lysed in 200 μL of 2XSDS reducing buffer, and lysates were heated at 95°C for 30 min followed by centrifugation at 15000 rpm for 15 min. Afterwards, 10 μL of cell lysate was loaded in each well and separated on SDS-PAGE followed by western blotting. The PVDF membranes were blocked in 5% BSA (in TBST) for 1 h followed by overnight incubation with phosphorylation site-specific p90RSK primary antibodies (phospho-Thr^359^, Cat. no. 8753, 1;500; phospho-Thr^573^, Cat. no. 9346S, 1:500; phospho-SerS^380^, CST, Cat. no. 11989, 1:500). Next day, blots were washed thrice with TBST for 10min each and incubated with anti-rabbit HRP-coupled secondary antibody (Genscript, Cat. no. A00098, 1:2000) for 1 h. The secondary antibody was rinsed off by washing the blots again with TBST for three times and developed with Promega ECL solution on chemidoc (BioRad). Blots were stripped with low pH stripping buffer and then re-probed for total RSK using RSK1/RSK2/RSK3 rabbit monoclonal primary antibody at 1:2500 dilution (CST, Cat. no. 9355S). Phospho-site specific signal was normalized with respect to the total RSK signal.

### Human monocyte-derived macrophages

Human monocyte-derived macrophages (HMDM) were derived and cultured following the previously described protocol (*21, 37*). Briefly, human buffy coat blood from anonymous healthy donors was obtained through the Australian Red Cross Blood Service (Brisbane, Australia). Human CD14^+^ monocytes were isolated from blood using Lymphoprep density centrifugation (STEMCELL, Melbourne, Australia) followed by CD14^+^ MACS separation (Miltenyi Biotec, Sydney, Australia). The isolated monocytes were cultured in Iscove’s Modified Dulbecco’s Medium (IMDM) containing 10% FBS, 100 U ml^−1^ penicillin, 100 μg ml^−1^ streptomycin and 15 ng ml^−1^ recombinant human macrophage colony stimulating factor (Lonza, Melbourne, Australia) on 10 mm square dishes (Bio-strategy, Brisbane, Australia). The adherent differentiated HMDMs were harvested by gentle scraping on Day 6-7.

### In-cell western assays on HMDMs

In-cell western assays were performed following the technical guidelines provided by LI-COR Biosciences (Lincoln, USA). Briefly, HMDMs were seeded (80,000 perwell) in poly D-lysine-coated (Merck, Perth, Australia) black-wall clear-bottom tissue culture 96-well plates (Corning, Corning, USA) for 24 h and serum-starved overnight. All ligands were prepared in serum-free IMDM containing 0.1% BSA (Merck, Perth, Australia). Cells were firstly pre-treated with the C5aR1 antagonist PMX53 (10 μM) for 20min (37 ℃, 5 % CO_2_) before stimulated with recombinant human C5a (Sino Biological, Beijing, China) or P32 (100 μM) for 10 min at room temperature. The media was removed and the cells were fixed using 4 % paraformaldehyde (Alfa Aesar, Haverhill, USA) (10min, room temperature). Upon gentle washing with DPBS, the cells were permeabilised using ice-cold methanol (10 min, room temperature) and then blocked using Odyssey Blocking Buffer in TBS (LI-COR Biosciences) (1.5h, room temperature). The cells were then stained with the indicated primary antibodies at 4℃ overnight (phospho-p90RSK^S380^, CST, Cat. no. 11989S, 1:800; phospho-p90RSK^T359^, CST, Cat. no. 8753S, 1:200; phospho-p90RSK^T573^, CST, Cat. no. 9346S, 1:200; Human/Mouse/Rat RSK Pan Specific Antibody, R&D Systems, Cat. no. RDSMAB2056, 1:200; phospho-p^44/42^ MAPK-ERK1/2^T202/Y204^, CST, Cat. no. 9101S, 1:250). Upon further washing with DPBS containing 0.1 % Tween-20, the cells were stained with IRDye 680RD donkey anti-rabbit secondary antibody (Cat. no. 926-68073, 1:1000,) and/or IRDye 800CW donkey anti-mouse IgG secondary antibody (Cat. no. 926-32212, 1:1000,) (LI-COR Biosciences, Lincoln, USA) for 1.5h at room temperature. The plate was then washed with DPBS containing 0.1 % Tween-20 and blotted dry. For fluorescence quantification, the plate was read on a Tecan Spark 20M microplate reader (Ex/Em: 667 nm/ 707 nm for IRDye 680RD and 770 nm/810 nm for IRDye 800CW, respectively) (Tecan, Männedorf, Switzerland).

### Neutrophil mobilization assay

Wild-type (WT), C5aR1−/− and C5aR2−/− mice on a C57BL/6J genetic background (n=5-15) were administered with recombinant mouse C5a (Sino Biological, China) at a dose of 50 μg kg-1 via intravenous injection (tail vein). After C5a injection, one drop of blood was collected from the tail tip to make a blood smear on a slide at 0, 15, 30 and 60 min. Blood smears were stained using a Microscopy Hemacolor^®^ Rapid Staining of Blood Smear Kit (Merck, Germany). Briefly, blood smears were fixed in Hemacolor^®^ Solution 1 (methanol). The slides were then stained with Hemacolor^®^ Solution 2 (Eosin Y), followed by Hemacolor^®^ Solution 3 (Azur B). The slides were washed with 1 x PBS (pH 7.2) and mounted with dibutyphthalate polystyrene xylene. Using a 20x/0.4 NA objective on an Olympus CX21 microscope, the first 400 white blood cells were counted for each slide, and the proportion of PMNs (i.e. cells containing granules that are light violet) was then calculated as previously described (*54*).

### Enzyme-linked immunosorbent assay

Blood for WT, C5aR1^−/−^ and C5aR2^−/−^ mice was collected in tubes containing 4 mM EDTA and plasma was obtained by centrifugation for 10 min at 2000xg at 4°C. Plasma TNFα levels were determined using a commercially available enzyme-linked immunosorbent assay kit (BD Biosciences).

### Data collection, processing, and statistical analysis

All the experiments were conducted at least three times and data were plotted and analyzed using GraphPad Prism software (Prism 8.0). Heat maps were plotted in Python 3.7 using appropriate libraries. The data were normalized with respect to proper experimental controls and appropriate statistical analyses were performed.

## Notes

### Competing Interest Statement

The authors have declared no competing interest.

