## Supplemental figures 1-10 for "Intrinsic bias at non-canonical, β-arrestin-coupled seven transmembrane receptors"

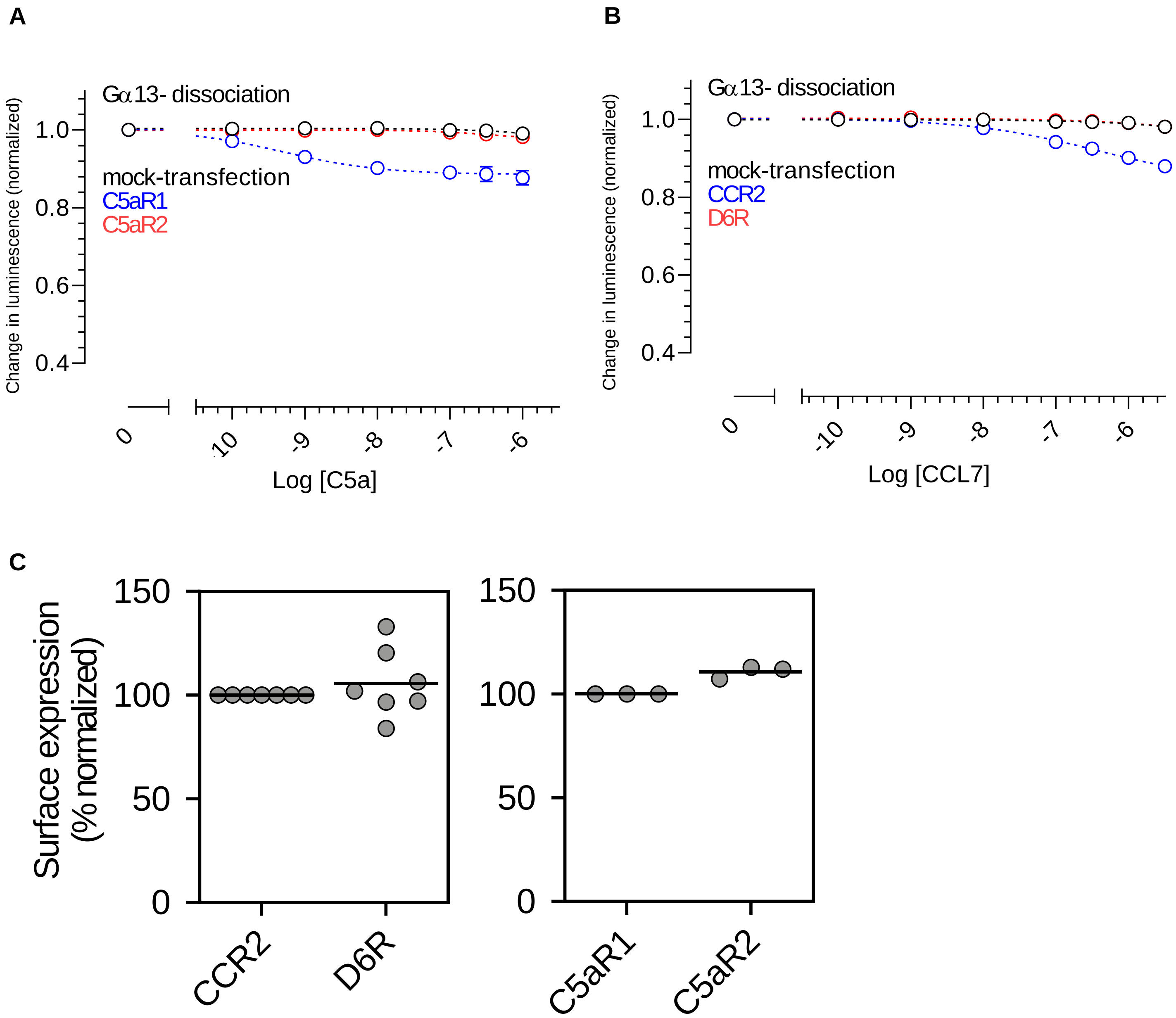

**Figure S1. Surface expression of the receptors and agonist-induced Gα13 activation. A-B.** Agonist-induced dissociation of Gα13 from Gβ1γ2 for C5aR1-C5aR2, and CCR2-D6R pairs measured using NanoBiT complementation assay. HEK-293 cells expressing the indicated receptor and Sm/Lg-BiT constructs of G-protein α,β,γ sub-units were stimulated with corresponding ligands, and the change in luminescence signal upon NanoBiT dissociation was measured as a readout of G-protein coupling and activation. Data represent three independent experiments, normalized with respect to baseline signal (i.e. before agonist-stimulation). **C.** Surface expression of the indicated receptors as measured in flow-cytometry based assay using an anti-FLAG tag monoclonal antibody and a secondary antibody conjugated with Alexa Fluor 488. Values of mean fluorescence intensity (MFI) from approximately 20,000 cells per sample were used for analysis.

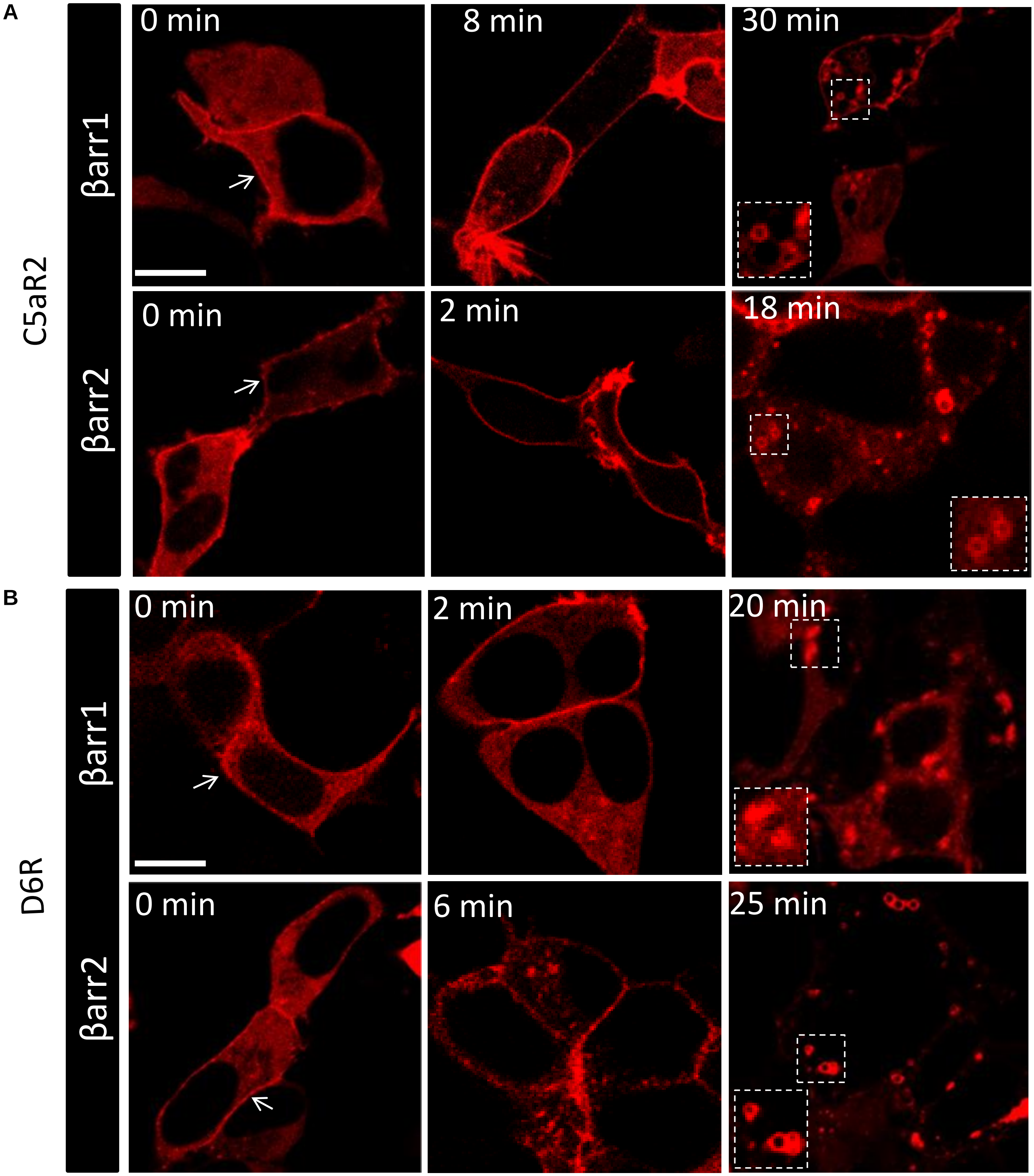

**Figure S2. Agonist-induced trafficking of mCherry-tagged βarr1/2 for C5aR2 and D6R.** HEK-293 cells expressing indicated receptors along with mCherry tagged βarr1 or βarr2 were observed for βarr trafficking in live cells upon specific agonist stimulation (D6R: CCL7 100nM; C5aR2: C5a 100nM). Both D6R and C5aR2 show constitutive βarr recruitment at the membrane under basal condition but trafficking to endosomes occurs only on ligand stimulation. There is apparent preference for βarr2 isoform over βarr1 in both D6R and C5aR2. Micrographs are representative images from three independent experiments at indicated time-points (scale bar, 10μm).

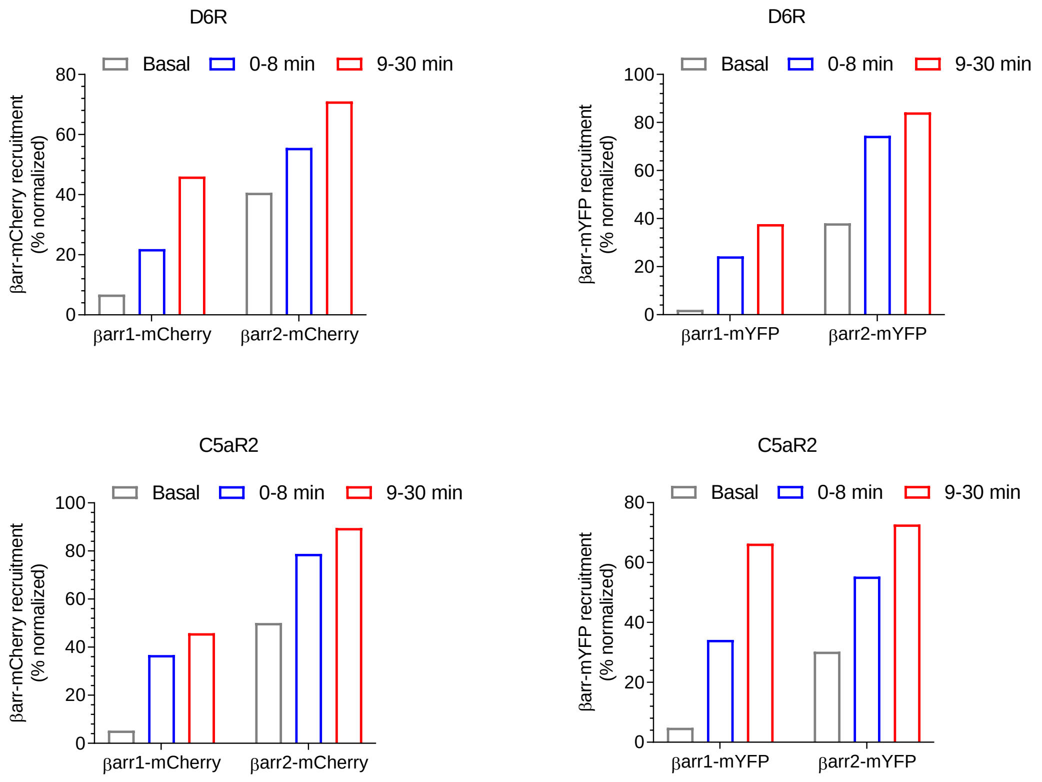

**Figure S3. Quantification of agonist-induced βarr translocation measured by confocal microscopy.** Agonist-induced localization of both mCherry and mYFP tagged βarr1 and 2 in D6R and C5aR2 were measured by confocal microscopy as described in material and methods section. βarr localization was manually scored in HEK293 cells from multiple fields from at least three independent experiments. Confocal images captured are grouped in three sections i.e. 0min, 1-8min and 9-30min post-agonist stimulation to reflect basal, early and late time-frames, respectively. The scoring of βarrs for localization at surface or endosome was done based on fluorescence signal from plasma membrane and punctate structures in the cytoplasm, respectively. Cells were scored from three independent experiments and data are plotted as % of βarr localization pattern from more than five hundred cells counted for each condition.

**A**

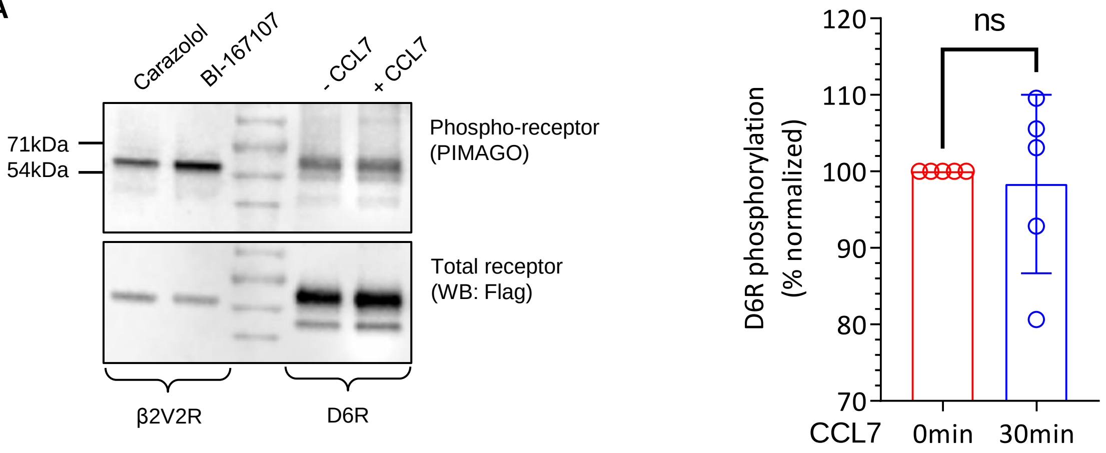

**B**

| | | |
|---|---|---|
| | Ser/Thr cluster | Distal C-terminus with acidic residues |
| D6R-C-terminus | AAVLGWHLAPGTAQASLSSCSESSILTAQEEMTGMNDLGERQSENYPNKEDVGNKSA | |
| D6R-Δ351 | AAVLGWHLAPGTAQASLSSCSESS | |
| D6R-Δ342 | AAVLGWHLAPGTAQA | |

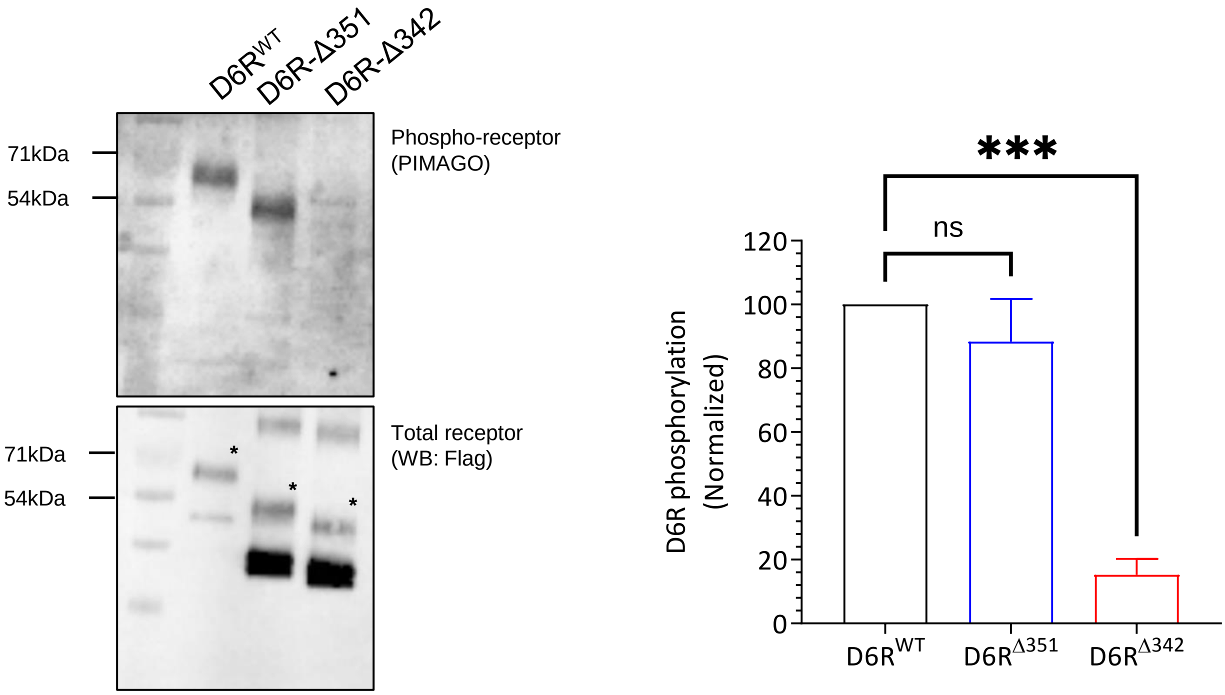

**Figure S4. D6R phosphorylation analysis. A.** D6R is constitutively phosphorylated as measured using PIMAGO kit, and its phosphorylation does not change upon ligand (CCL7) stimulation (n=5). As a control, a chimeric β2-adrenergic receptor construct was used, which exhibits agonist-induced phosphorylation**.** A part of this data is presented as Figure 3C. **B.** The primary phosphorylation determinants of D6R are localized in the region between Ser$^{351}$ and Ser$^{342}$ as the receptor truncation at Ser$^{342}$ but not at Ser$^{351}$ abolishes constitutive phosphorylation. All three constructs were expressed at comparable levels in HEK-293 cells, and only the matured receptor population (indicated with an asterisk on the anti-Flag blot) was used for normalization (n=3).

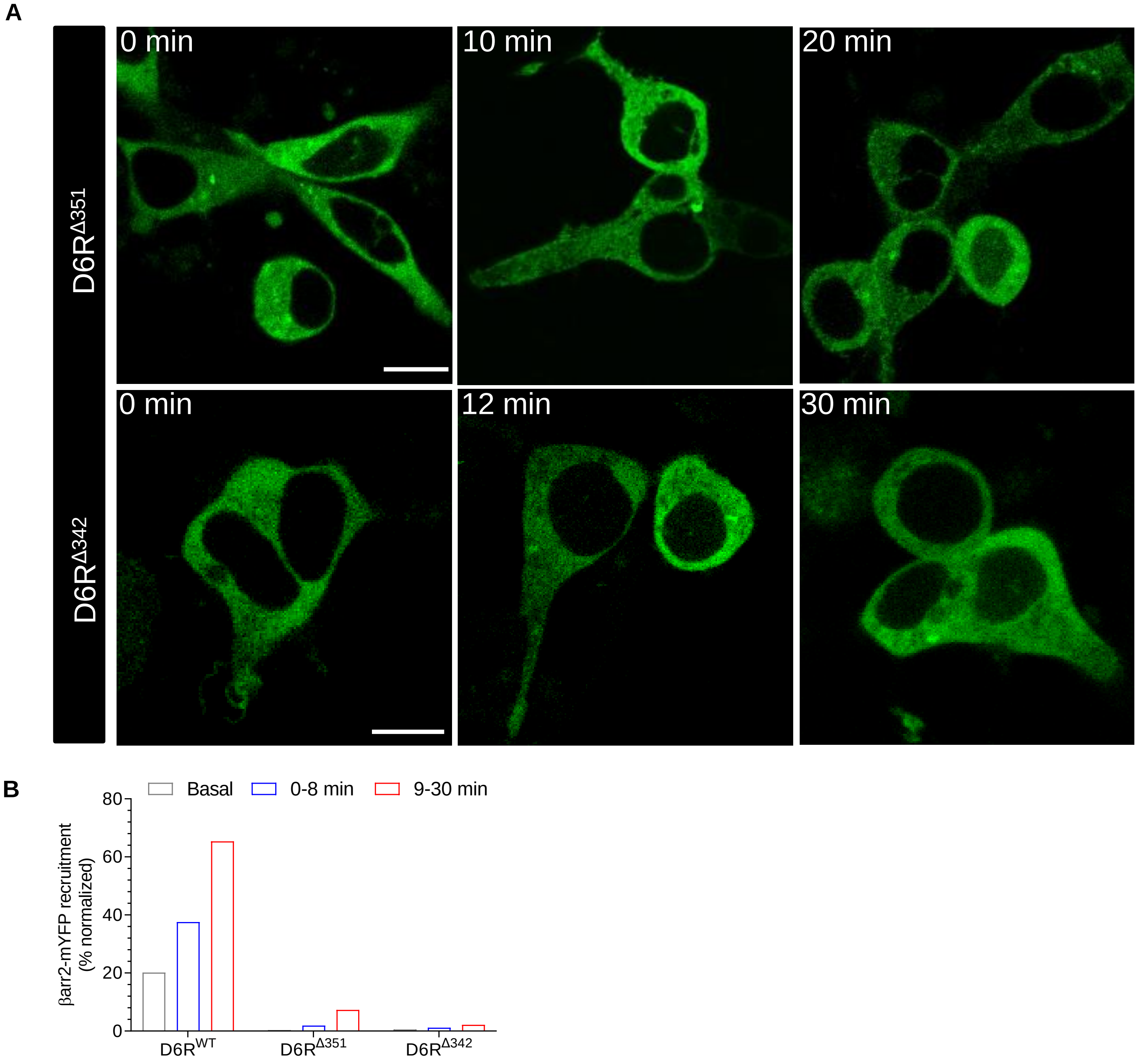

**Figure S5. Effect of carboxyl-terminus truncation on agonist-induced βarr recruitment. A.** HEK-293 cells expressing the indicated receptor constructs and βarr2-mYFP were stimulated with agonist (100nM CCL7) and localization of βarr2 was monitored by confocal microscopy (Scale bar is 10μm). **B.** For assessing βarr recruitment and trafficking in D6R deletion mutants, βarr2 localization was manually scored in HEK293 cells from multiple fields from at least three independent experiments. Confocal images captured are grouped in three sections i.e. 0min, 1-8min and 9-30min post-agonist stimulation to reflect basal, early and late time-frames, respectively.

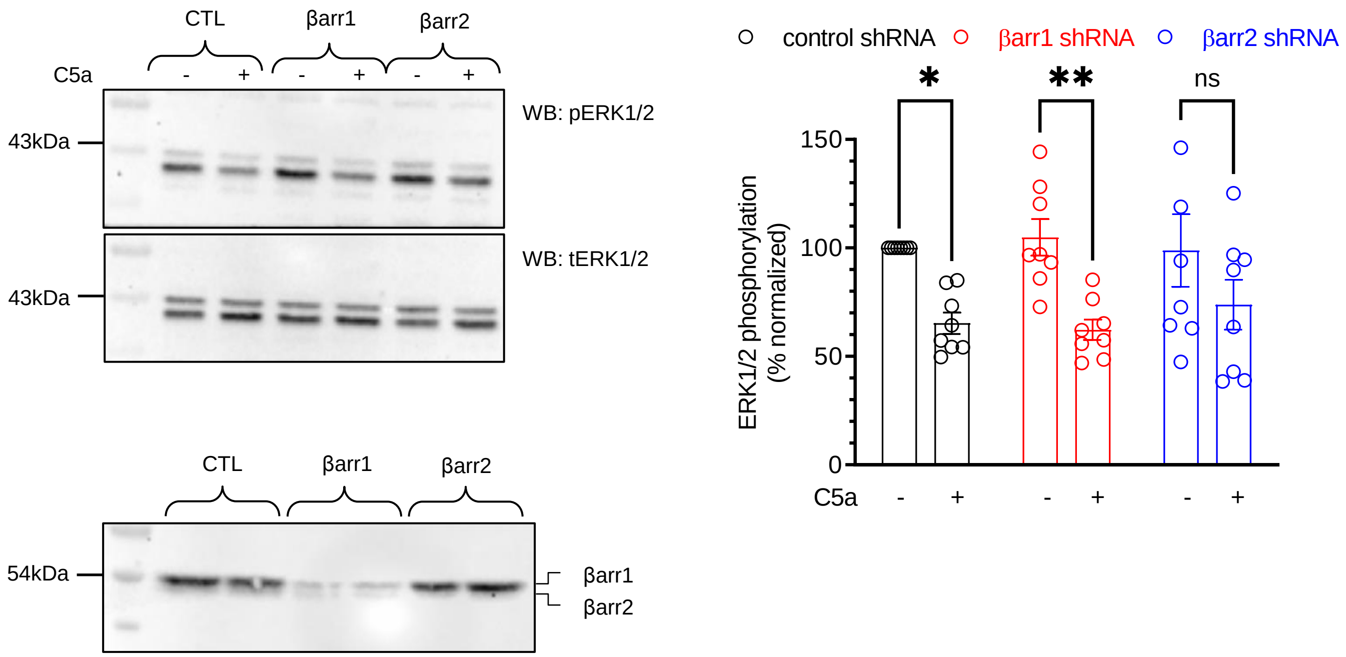

**Figure S6**. **Effect of βarr depletion on ERK1/2 phosphorylation**. C5aR2-expressing cells exhibit an elevated level of pERK1/2, which is reduced upon C5a-stimulation. shRNA-mediated depletion of βarr2 but not βarr1 attenuates the C5a-induced lowering of pERK1/2 while knock-down of either βarrs have no significant effect on basal pERK1/2. The left panel shows a representative image from three independent experiments and the right panel show densitometry-based quantification from eight independent experiments normalized with respect to the basal pERK1/2 under control shRNA conditions (* $p<0.05$; ** $p<0.01$). The lower panel shows shRNA mediated depletion of βarrs in HEK-293 cells used in these experiments.

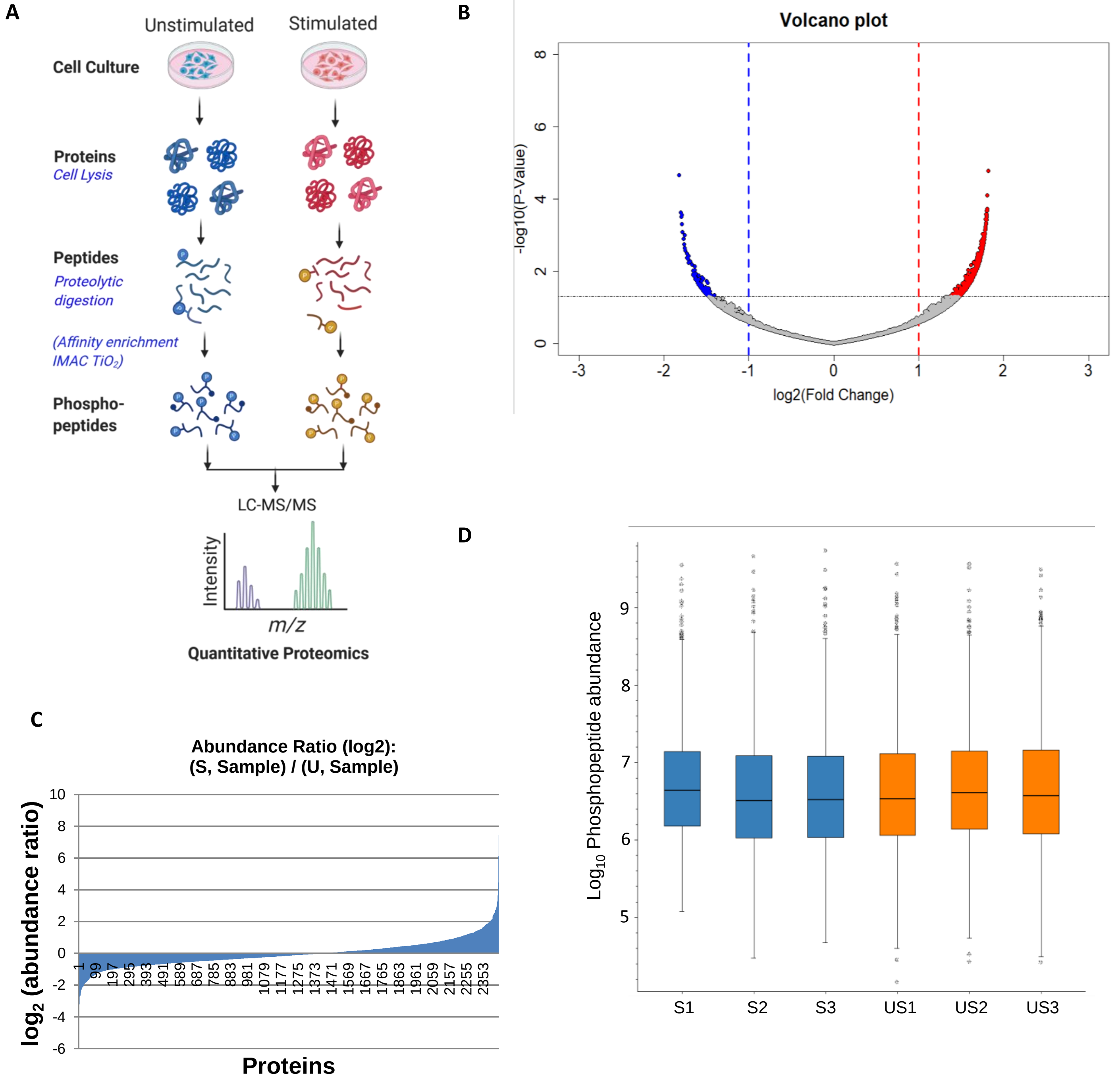

**Figure S7. Phosphoproteomics analysis on D6R expressing cells upon CCL7-stimulation. A.** Schematic representation of the workflow for sample preparation and LC-MS/MS based phospho-proteomics study. **B.** Volcano plot showing the difference in log fold change of phosphopeptides in unstimulated compared to stimulated samples. P-values of t-test shows significant difference in abundance of phospho-up regulated (represented in red) and phospho-down regulated (represented in blue) phosphopeptides across three sets of independent biological replicates. **C.** Differential expression of 1132 proteins reflected by the abundance ratio (log2) in stimulated vs unstimulated samples. **D.** Boxplots of three different biological replicates from stimulated and unstimulated samples from six different runs showing comparatively similar features among different samples (S1,S2,S3 = stimulated; US1,US2,US3 = unstimulated).

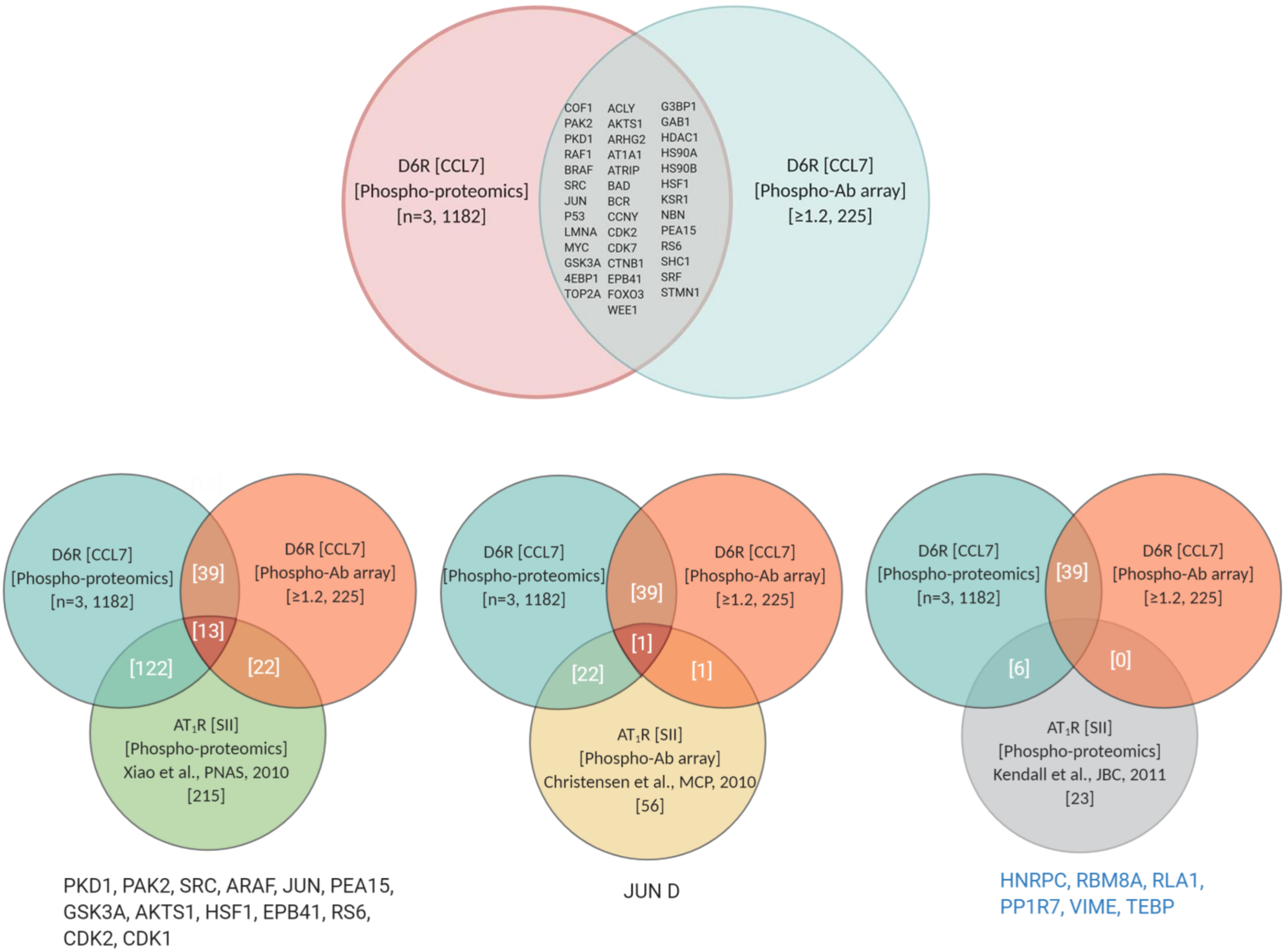

**Figure S8. Analysis of phospho-antibody array and phosphoproteomics hits.** The upper panel shows a Venn diagram comparing the protein identified in the phosphoproteomics and phospho-antibody array screens performed on D6R expressing cells. The common proteins are indicated in the overlapping region with the generic gene symbols. The lower panels show the Venn diagram comparing the D6R phospho-hits with previous studies carried out on AT1R stimulated with a βarr-biased ligand, SII. The common hits identified in this analysis are listed and indicated by their gene symbol.

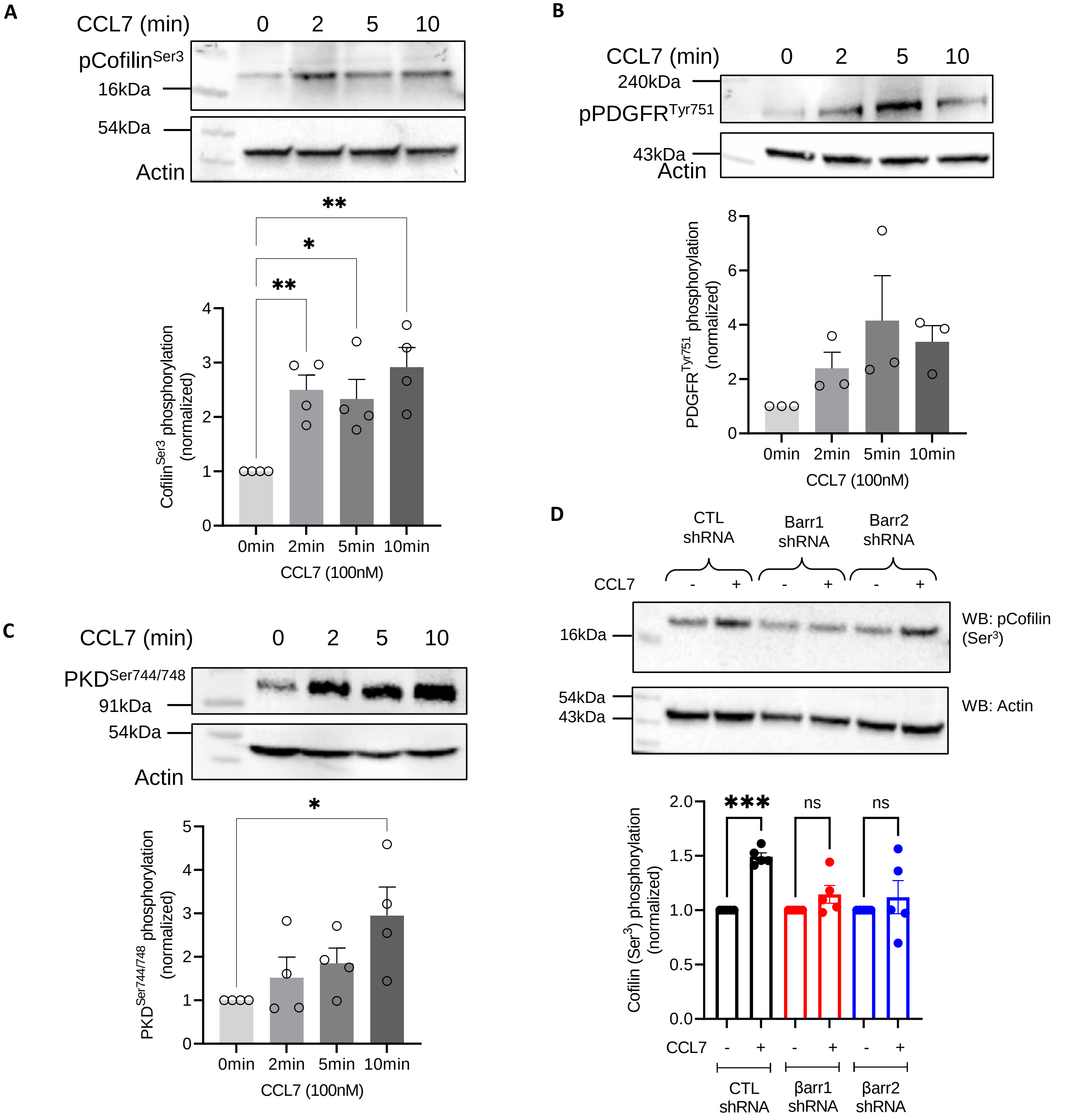

**Figure S9. Validation of D6R phospho-hits in HEK-293 cells. A-C.** Phosphorylation of cofilin, PDGFR-β and PKD1 upon CCL7-stimulation of D6R in HEK-293 cells was validated using Western blotting with corresponding antibodies. A representative blot from 3-5 experiments and densitometry-based quantification (average±sem) is presented (*p<0.05, ***p<0.001; One-Way ANOVA). **D.** shRNA-mediated depletion of βarr1 and βarr2 and attenuate CCL7-induced (200nM) phosphorylation of cofilin (Ser³). The upper panel shows a representative image from three independent experiments and the lower panel shows densitometry-based quantification, normalized with respect to the basal coffilin phosphorylation (i.e. without CCL7-stimulation). Data are analyzed using One-Way ANOVA; **p<0.01).

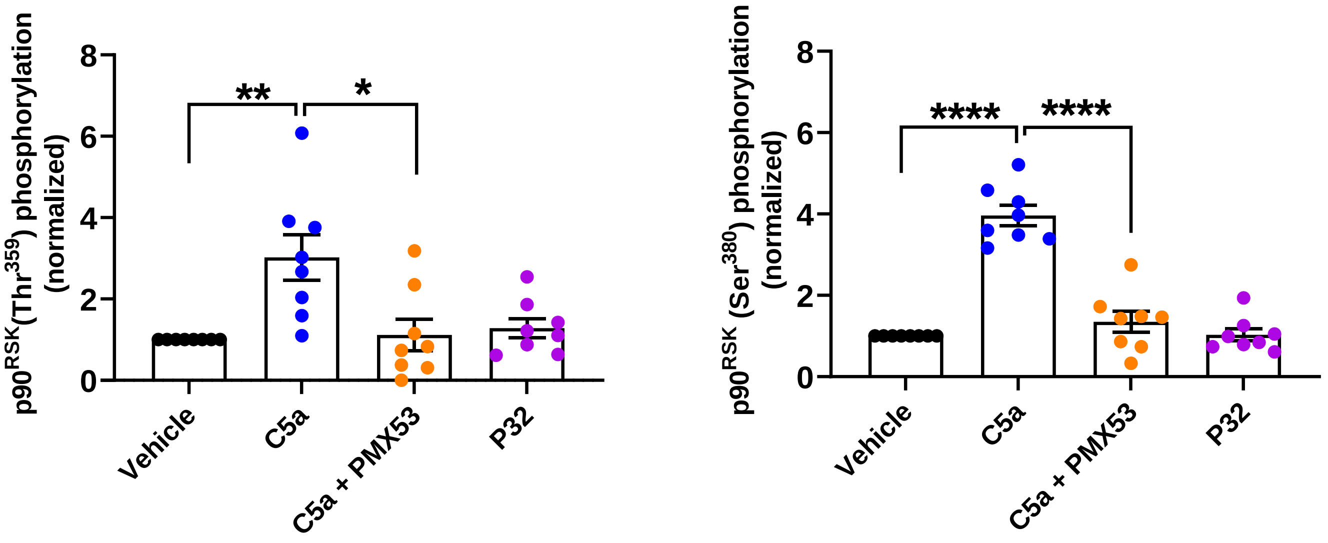

**Figure S10**. **Agonist-induced p90$^{RSK}$ phosphorylation in HMDMs.** The phosphorylation of p90$^{RSK}$ at Thr$^{359}$ and Ser$^{380}$ are primarily mediated through C5aR1 activation as pre-treatment with PMX53 (C5aR1 antagonist) blocks C5a-induced response. Moreover, P32 (C5aR2-specific agonist) stimulation of HMDMs also fails to induce significant phosphorylation on these sites. Data from HMDMs derived from eight different donors were collected and normalized with respect to vehicle-treatment, and analyzed using Fisher's LSD test, assuming each treatment group is independent of each other - only donors with a >1 response to C5a are included.
